# Methionine Metabolism Controls the B-cell EBV Epigenome and Viral Latency

**DOI:** 10.1101/2022.02.24.481783

**Authors:** Rui Guo, Jin Hua Liang, Yuchen Zhang, Michael Lutchenkov, Zhixuan Li, Yin Wang, Vicenta Trujillo-Alonso, Rishi Puri, Lisa Giulino-Roth, Benjamin E. Gewurz

## Abstract

Epstein-Barr virus (EBV) subverts host epigenetic pathways to switch between viral latency programs, colonize the B-cell compartment and reactivate. Within memory B-cells, the reservoir for lifelong infection, EBV genomic DNA and histone methylation marks restrict gene expression. But, this epigenetic strategy also enables EBV-infected tumors, including Burkitt lymphomas to evade immune detection. Little is known about host cell metabolic pathways that support EBV epigenome landscapes. We therefore used amino acid restriction, metabolomic and CRISPR approaches to identify that an abundant methionine supply, and interconnecting methionine and folate cycles, maintain Burkitt EBV gene silencing. Methionine restriction, or methionine cycle perturbation, hypomethylated EBV genomes, de-repressed latent membrane protein and lytic gene expression. Methionine metabolism also shaped EBV latency gene regulation required for B-cell immortalization. Dietary methionine restriction altered murine Burkitt xenograft metabolomes and de-repressed EBV immunogens *in vivo*. These results highlight epigenetic/immunometabolism crosstalk supporting the EBV B-cell lifecycle and suggest therapeutic approaches.

**Highlights:**

- Methionine metabolism is critical for Epstein-Barr virus B-cell latency
- Extensive cross-talk enables methionine metabolism to control the EBV epigenome
- Methionine restriction also impairs EBV-driven human B-cell immortalization
- Dietary methionine restriction unmasks EBV antigens in Burkitt xenografts in vivo

## Introduction

A hallmark of the Epstein-Barr virus (EBV) lifecycle is the use of multiple viral genome programs to navigate the B-cell compartment in order to colonize the memory B-cell reservoir (Buschle and Hammerschmidt, 2020; Guo and Gewurz, 2021; Pei and Robertson, 2020). In order to persistently infect B-cells, which serve as professional antigen presenting cells at the heart of the adaptive immune system, EBV must therefore carefully regulate when and where it expresses 80 virally-encoded protein antigens. This strategy enables EBV to infect approximately 1 in 100,000 memory B-cells and to periodically undergo lytic reactivation, with expression of all EBV proteins, even in immunocompetent hosts. Much remains to be learned about how EBV exploits crosstalk between host immunometabolism and epigenetic pathways in order to regulate eight viral latency and nearly 80 lytic cycle programs to manipulate B-cell biology and evade immunity.

EBV persistently infects 95% of adults worldwide. While EBV typically establishes a balance with the host, acute infection can result in infectious mononucleosis, a syndrome characterized by polyclonal B-cell activation and potent anti-EBV immune responses. EBV is increasingly appreciated to be a major trigger for multiple sclerosis and other autoimmune diseases (Bjornevik et al., 2022; Laderach and Munz, 2021; Lanz et al., 2022). EBV is also associated with 200,000 cancer per year, including Burkitt, Hodgkin, post-transplant and T/NK cell lymphomas, as well as gastric and nasopharyngeal carcinoma (Farrell, 2019; Gewurz et al., 2021; Shannon-Lowe et al., 2017). Each of these cancers exhibits a characteristic EBV latency program, in which epigenetic mechanisms dictate the combination of EBV genes that are expressed and suppress the production of infectious virion (Price and Luftig, 2015). Patterns observed in these human malignancies mirror EBV latency program expression at distinct stages of B-cell activation and differentiation.

Upon B-cell infection, the EBV double stranded DNA genome is chromatinized and modified by DNA and histone epigenetic marks. Together with three dimensional genome architecture, these culminate in the activation of the viral C promoter, which drives the latency III program, comprised of six Epstein-Barr nuclear antigens (EBNA), two latent membrane protein (LMP) oncoproteins and non-coding RNAs (Tempera and Lieberman, 2014). Latency III is observed in post-transplant and central nervous system lymphomas. EBNA2 is a major activator of the LMP promoters in latency III and serves as a master regulator of host metabolic pathways (Alfieri et al., 1991; Gewurz et al., 2021). EBNA2 highly induces one-carbon and lipid metabolism pathways (Wang et al., 2019b; Wang et al., 2019c), and is essential for the outgrowth of newly EBV-infected primary human B-cells over the first 8 days of infection (Pich et al., 2019). Epigenetic marks also silence the immediate early gene BZLF1 to prevent expression of the nearly 80 EBV lytic genes (Buschle and Hammerschmidt, 2020; Chakravorty et al., 2019).

As infected cells enter secondary lymphoid tissue germinal centers, epigenetic mechanisms switch the EBV genomic program to latency II (Guo and Gewurz, 2021; Price and Luftig, 2015). In this state, repressive DNA methylation marks silence the C promoter to down-modulate expression of five EBNA antigens. In the absence of C promoter activity, the genome tethering EBNA1 is expressed from the viral Q promoter, whereas undefined mechanisms sustain LMP promoter activity. LMP1 and LMP2A mimic signaling from CD40 and B-cell receptors, respectively, driving B-cell growth and survival pathways. Hodgkin lymphoma Reed-Sternberg cells, which originate from germinal centers, exhibit the latency II program (Alfieri et al., 1991; Gewurz et al., 2021). Little is presently known about how immunometabolism pathways cross-talk with the EBV epigenome in the transition to latency II.

EBV-infected cells differentiate into memory B-cells, where epigenetic mechanisms also silence LMP1 and LMP2A expression. In this latency I program, EBNA1 is the only EBV-encoded protein expressed (Buschle and Hammerschmidt, 2020; Chakravorty et al., 2019; Gewurz et al., 2021; Guo and Gewurz, 2021). This property enables EBV+ memory cells to evade immune detection, a property shared by Burkitt lymphoma B cells that also typically exhibit the latency I program. DNA methylation is critical mechanism by which EBNA and LMP expression is silenced in Burkitt tumor cells and presumably also in memory cells. The epigenetic writer polycomb repressive complex II has an additional role in silencing the LMP1 and LMP2A promoters in Burkitt cells (Guo et al., 2020b).

EBV highly remodels newly infected B-cell metabolic pathways (Bonglack et al., 2021; Lamontagne et al., 2021; McFadden et al., 2016; Mrozek-Gorska et al., 2019; Wang et al., 2019b; Wang et al., 2019c). Amino acid transporters, including those necessary for methionine import, are amongst the most highly EBV upregulated at the plasma membrane (Wang et al., 2019b). Despite this signal, nutritional effects on the EBV/host relationship remains largely undefined. Likewise, the metabolism master regulator MYC is highly expressed in Burkitt tumors, which increases levels of methionine import and metabolism (Dang, 2016; Yue et al., 2017). Methionine cycle metabolism produces the universal methyl donor S-adenosylmethionine (SAM), which supplies DNA and histone methyl transferases to drive epigenetic modifications. The byproduct S-adenosylhomocysteine (SAH) competitively inhibits methyltransferase reactions (Dai et al., 2020; Ducker and Rabinowitz, 2017; Sanderson et al., 2019). Therefore, the SAM:SAH ratio serves as an indicator of the cellular methylation potential.

Methionine cycle flux converts SAH to homocysteine, which is further metabolized to regenerate methionine upon donation of a carbon unit from interconnecting folate metabolism pathways (Herbig et al., 2002; Lu and Mato, 2012). EBNA2 and its key host oncoprotein target MYC highly upregulate EBV-infected B-cell folate and one carbon metabolism pathways, at least in part to support de novo nucleotide synthesis and redox defense (Wang et al., 2019b). How interconnecting methionine and folate metabolism networks cross-talk with the viral and host epigenomes has yet to be studied in the context of EBV infection. Mammals exclusively obtain methionine from dietary sources, and restriction of dietary methionine uptake can strongly impact both the availability of methyl donors and metabolic disease hallmarks (Dai et al., 2018; Maddocks et al., 2016; Quinlan et al., 2017; Shiraki et al., 2014). Whether methionine restriction can impact the EBV epigenome remains undefined.

In this study, we investigate methionine and folate metabolism pathway roles in EBV epigenome regulation in Burkitt and newly infected B-cells. We identify obligatory methionine and folate metabolism roles in viral genome histone and DNA methylation necessary for suppression of the EBV latency III and lytic antigens. Conversely, primary human B-cell infection assays demonstrate obligatory methionine metabolism roles in EBV oncoprotein expression programs and B-cell growth transformation, a model for post-transplant lymphoproliferative disease. Collectively, these studies highlight multiple therapeutic targets for EBV latency reversal approaches, including dietary methionine restriction.

## Results

### Methionine restriction de-represses EBV lytic and latent genes

The extent to which Burkitt cells require high flux through methionine uptake pathways is unknown. Methionine serum concentrations normally range from 10-30 μM, but can be significantly impacted by diet (Schmidt et al., 2016). By comparison, the methionine concentration in Roswell Park Medical Institute 1640 (RPMI) medium commonly used for *in vitro* B-cell culture is ∼100 μM. To gain insights into methionine metabolism roles in control of the EBV epigenome and transcriptome, we investigated the effects of methionine restriction (MR) on a panel of Burkitt cells with well-defined EBV latency programs. We first tested MR effects on Mutu I cells, which were established from a latency I African BL tumor (Gregory et al., 1990). Intriguingly, culture for 72 hours in RPMI with 10 μM methionine induced LMP1 and LMP2A (Figure 1A**, S1A**), whose expression is typically repressed by DNA methylation in EBV latency I (Falk et al., 1998; Guo et al., 2020b; Tao et al., 1998). Unexpectedly, MR failed to de-repress EBNA2, and only modestly induced EBNA3 expression (**Figure S1B**), perhaps indicating a lower threshold for de-repression of the LMP promoter. We therefore next tested MR effects on P3HR-1 cells, in which a genomic deletion knocks out EBNA2, the major viral activator of the LMP promoter. Despite the absence of EBNA2, MR highly de-repressed LMP1 and LMP2 expression (Figure 1B), suggesting that methionine metabolism is required for silencing of the LMP1 promoter independently of effects on EBNA2.

**Figure 1.**
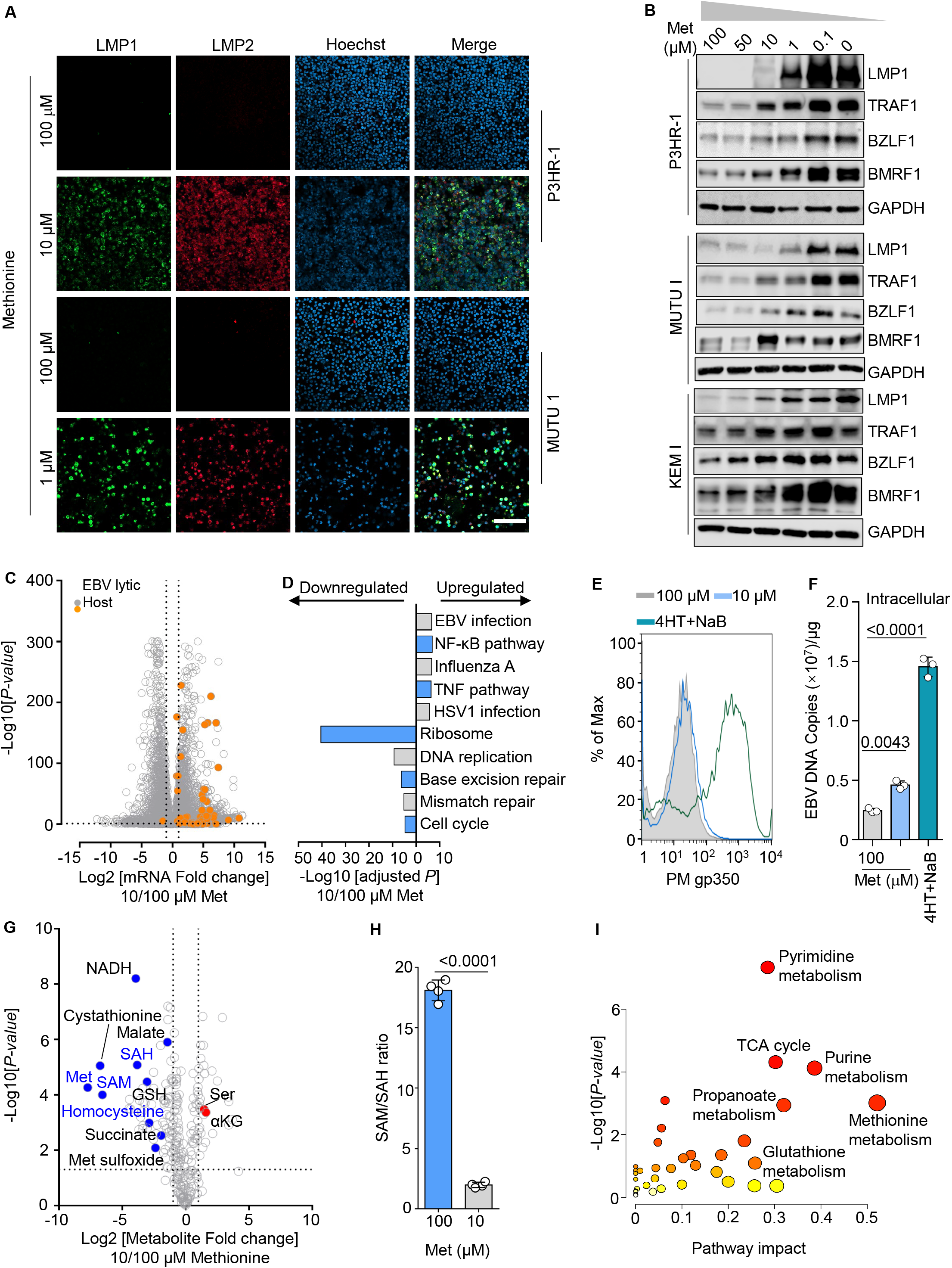
Methionine restriction de-represses Burkitt latent membrane and lytic antigens. A) Confocal immunofluorescence analysis of LMP1 or LMP2A expression in P3HR-1 or Mutu I cells cultured in media with the indicated methionine concentrations for 3 days. Scale bar, 100 µm. B) Immunoblot analysis of whole cell lysates (WCL) from P3HR-1, Mutu I, or KEM I cells cultured in media with indicated concentration of methionine (Met) for 3 days, representative of n=3 experiments. C) Volcano plot visualization of −Log10 (p-value) statistical significance (y-axis) and Log2 fold-change in mRNA abundance (x-axis) from RNAseq analysis of P3HR-1 cells cultured in 10 vs. 100 μM Met for 3 days. Data are from duplicates RNA-seq datasets. EBV (orange) and host (gray) values are shown. D) Enrichr KEGG pathway –Log 10 (adjusted p-values) of gene sets significantly changed in 100 versus 10 μM Met, as in (D). E) FACS analysis of plasma membrane (PM) gp350 expression in P3HR-1 ZHT/RHT cells grown with 100 versus 10 μM Met for 3 days or reactivated with 4HT (1µM) and sodium butyrate (NaB, 3mM) as a positive control. F) qRT-PCR analysis of EBV intracellular genome copy number from cells described in (E). Total genomic DNA was extracted at 72h in the indicated media or at 48h post 4HT/NaB. Mean ± SD values from n=3 replicates are shown. G) Volcano plot of metabolomic analysis of P3HR-1 cells grown in 100 versus 10 μM Met for 72 hours. Methionine cycle metabolites are highlighted in blue text. H) SAM/SAH ratio from cells treated as in (G). Mean ± SD values from n=4 replicates are shown. I) Metabolic pathway analysis plot showing the most strongly impacted metabolic pathways downregulated by MR. The x-axis represents pathway impact value computed from MetaboAnalyst 3.0 topological analysis, y-axis is the-log of the P-value obtained from pathway enrichment analysis. See also Figures S1-3.

To gain further insights into MR effects on EBV genome reprogramming, we next analyzed Burkitt cell responses to a wider range of methionine concentrations. LMP1 de-repression, and induction of its downstream transcriptional target TRAF1 (Zhao et al., 2014), were observed in all three cell lines, indicating activation of LMP1/NF-κB pathways in a dose-responsive manner. Similarly, MR induced plasma membrane expression of LMP1/NF-κB target ICAM-1 in all five EBV+ Burkitt cell lines tested, but not in a closely matched EBV negative Akata cell line or in EBV negative REH cells (**Figure S1C**). Since DNA methylation is also necessary for Burkitt cell repression of EBV lytic genes, we also analyzed immediate early BZLF1 and early BMRF1 gene expression. Both BZLF1 and BMRF1 were de-repressed by MR in a dose-responsive manner (Figure 1B), suggesting obligatory methionine metabolism roles in Burkitt latency maintenance. Flow cytometry (FACS) analysis of 7-Aminoactinomycin D (7-AAD) vital dye uptake indicated little effect on BL viability at day 3 post MR, while CFSE dye-dilution and cell cycle analyses showed that MR caused a G1 growth arrest, despite EBV oncoprotein induction (**Figure S1D-F**).

To more broadly define MR effects on EBV and host gene expression, RNAseq was performed in P3HR-1 cells grown in 100 vs 10 μM methionine for 72 hours. MR significantly depressed 60 EBV lytic genes as well as LMP1, 2A and EBNA3 transcripts (Figure 1C**, S2 and Supplementary Tables S1-2**). Thus, MR may perturb EBNA3 expression at the protein level, given apparently greater increases in EBNA3 mRNA than protein levels. Highly concordant effects were observed in Rael Burkitt cells, in which DNA methylation enforces as tight latency I phenotype (Dalton et al., 2020; Ernberg et al., 1989; Guo et al., 2020b; Masucci et al., 1989) (**Figure S2A-B**). Notably, even though Rael EBV genomes have an intact EBNA2 locus, EBNA2 mRNA abundance was not significantly increased by MR, even though MR increased EBNA3, LMP and lytic gene expression. EBNA-LP mRNA abundance was also only modestly increased, suggesting a methionine metabolism role in control of the mRNA encoding EBNA2 and LP (**Figure S2A**). Host gene ontology analysis identified that MR significantly upregulated NF-κB pathway targets, suggestive of signaling by de-repressed LMP1 (Figure 1D**, S2C**). RNAs encoding ribosomal components were strikingly downregulated in both P3HR-1 and Rael cells (Figure 1D**, S2C**), indicating unexpectedly strong cross-talk between methionine and ribosomal metabolism in Burkitt cells. qPCR analysis validated effects on Rael cell immediate early BZLF1, early BMRF1 and late BLLF1 induction (**Figure S2D**).

Despite immediate early and early gene expression, MR only weakly induced EBV late gene expression (**Supplementary Tables S1-2**). To further examine this result, we performed FACS analysis of P3HR-1 cells grown in 10 vs 100 μM methionine. MR did not significantly increase late gene gp350 plasma membrane abundance, in contrast with a full lytic cycle induced by conditional BZLF1 expression and sodium butyrate (Figure 1E). We therefore asked whether MR induced EBV lytic DNA replication, which is necessary for late gene expression. Culturing P3HR-1 cells with 10 μM methionine only modestly increased EBV DNA copy number, in comparison with positive control activation of a conditional BZLF1 allele together with HDAC inhibitor sodium butyrate treatment (Figure 1F). Taken together, these results indicate that MR depresses immediate early and early genes, but generally results in an abortive lytic cycle.

MR induction of LMP1 and LMP2A, in the absence of EBNA2, raised the question of whether EBV lytic reactivation was responsible for their expression. In support, LMP1 is induced in both epithelial and B-cells upon lytic reactivation (Webster-Cyriaque et al., 2000; Yuan et al., 2006). To test this hypothesis, we generated CRISPR BZLF1 knockout Mutu I cells to prevent lytic reactivation, and as a control for EBV episome CRISPR editing, also generated Mutu I knocked out for the late gene BLLF1/gp350. We confirmed that BZLF1 KO prevented induction of the early BMRF1 antigen in response to immunoglobulin-crosslinking (**Figure S2E**). Despite being unable to trigger lytic gene expression as evidenced by lack of BMRF1 induction, MR strongly induced LMP1 in BZLF1 KO cells, suggesting latent cycle LMP promoter activation by a host transcription factor reminiscent of EBV latency II (**Figure S2F**). These results indicate that MR independently blocks silencing of EBV lytic and LMP1 antigens.

To gain insights into how MR alters Burkitt cell metabolic pathways that result in EBV gene de-repression, we performed liquid chromatography mass spectrometry (LC/MS) metabolomic analyses of P3HR-1 cells grown with 100 vs 10 μM for 72 hours. As expected, the abundance of methionine was amongst the most downregulated by MR, as were the methionine cycle metabolites homocysteine, SAM and SAH. Importantly, intracellular SAM and SAH were reduced about 223 and 11-fold, respectively, indicating that the cellular methylation potential was decreased by ∼9-fold (Figure 1G-H**, S3A**). Notably, MR upregulated the abundance of α-ketoglutarate, which serves as a necessary cofactor for TET family DNA demethylases, but decreased levels of succinate, which instead inhibits TET demethylase activity (Figure 1G-H). MR also downregulated metabolites of Burkitt purine and pyrimidine metabolism, TCA and transsulfuration pathways (Figure 1I). Transsulfuration uses methionine for *de novo* cysteine synthesis. Collectively, these results highlight that the Burkitt B-cell metabolome is highly influenced by extracellular methionine abundance, and that methionine metabolism exerts key effects on the maintenance of the highly restricted EBV latency I program in Burkitt cells.

### MR alters the EBV epigenome and reprograms the host epigenetic factor landscape

Given MR effects on the cellular methylation potential, we next investigated how MR alters the EBV-infected cell DNA and histone methylation epigenetic landscapes. 5-methylcytosine (5mC) dot blot analysis identified significant decreases in global DNA methylation in methionine restricted cells (Figure 2A). Similarly, immunoblot for histone methylation epigenetic marks demonstrated reduced histone 3 lysine 9 tri-methyl (H3K9me3) and histone 3 lysine 4 tri-methyl H3K4me3 marks. However, repressive histone 3 lysine 27 tri-methyl (H3K27me3) were not significantly changed by 72 hours of MR (Figure 2B), indicating that histone epigenetic marks were not universally diminished. These results contrast with recent T-cell analyses, where MR reduced H3K4me3 and H3K27me3 but not H3K9me3, suggesting a degree of cell type specificity (Roy et al., 2020).

**Figure 2.**
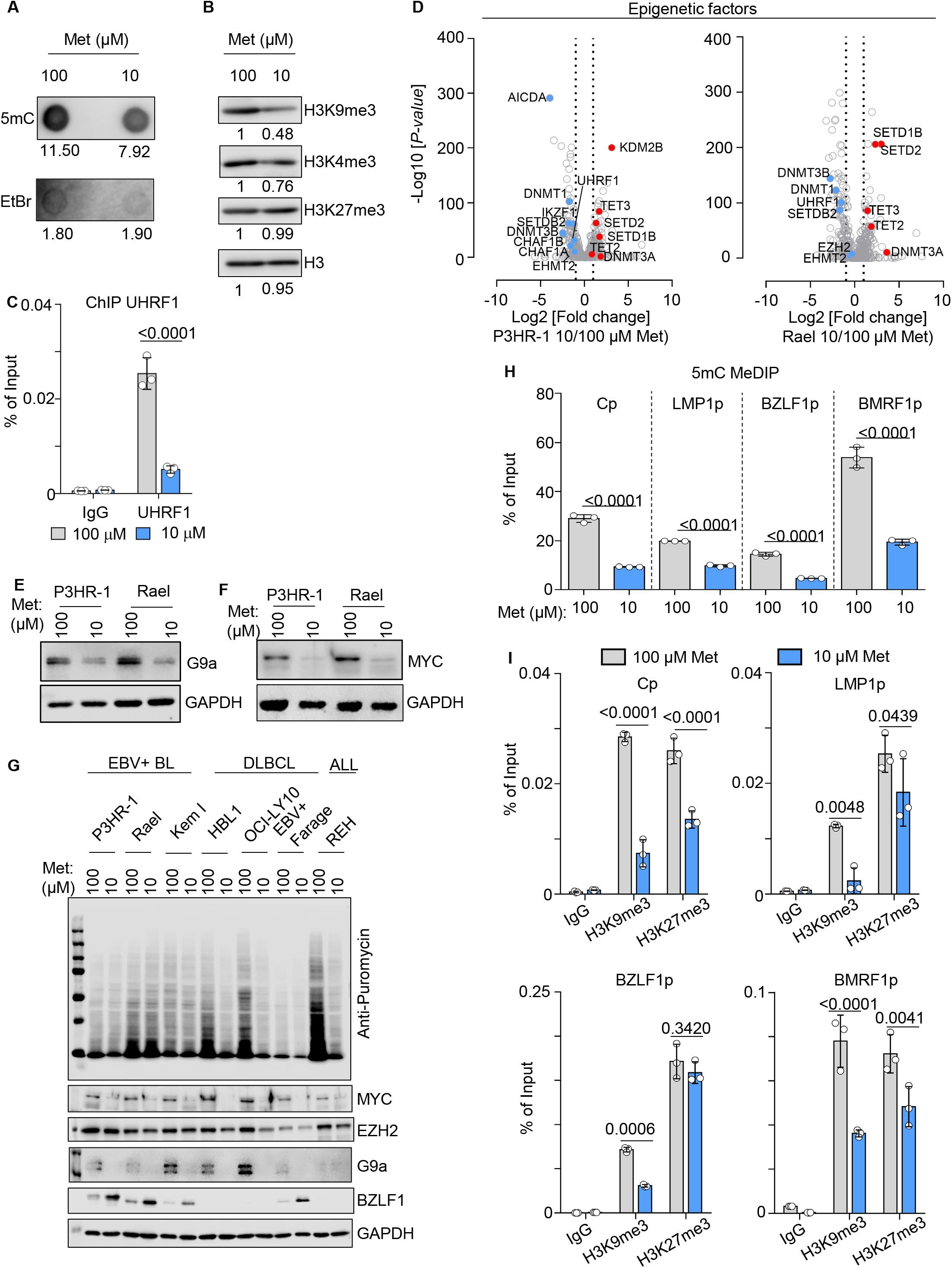
Methionine restriction alters the host and EBV epigenetic landscape. A) 5mC dot blot analysis of DNA extracted from P3HR-1 cultured in 100 vs 10 μM Met for 72h. As a loading control, membranes were stained with ethidium bromide. B) Immunoblot analysis of WCL from P3HR-1 cells cultured in cultured in 100 vs 10 μM Met for 72h. C) ChIP-qPCR analysis of IgG control versus anti-UHRF1 abundances in chromatin extracted from P3HR-1 cells cultured in 100 vs 10 μM Met for 72h. Shown are *Cp* qPCR mean ± SD values from n=3 replicates. D) Volcano plot of host epigenetic factor RNAseq from P3HR-1 (left) or Rael (right) cells cultured in 100 vs 10 μM Met for 72h, using the curated EpiFactors database (Medvedeva et al., 2015). P value and log fold change data were generated from n=2 replicates (P3HR-1) or n=3 replicates Rael by DESeq2, using default settings with Wald test and normal shrinkage, respectively. E-F) Immunoblot analysis of WCL from P3HR-1 or Rael cultured in 100 vs 10 μM Met for 72h. G) Immunoblot analysis of puromycin incorporation into newly synthesized polypeptides (top) or of the indicated proteins from WCL prepared from the indicated B cell lines cultured in 100 vs 10 μM Met for 72h. H) 5mC MeDIP analysis of chromatin from P3HR-1 cultured in 100 vs 10 μM Met for 72h. Shown are mean ± SD values from n=3 replicates of qPCR analysis of the C, LMP1, BZLF1 or BMRF1 promoters. Cells were maintained in acyclovir (100µg/ml) to prevent lytic DNA synthesis. I) ChIP for H3K9me3 or H3K27me3 of chromatin from P3HR-1 cells cultured in 100 vs 10 μM Met for 72h, followed by qPCR with primers specific for the C, LMP1, BZLF1 or BMRF1 promoters. Cells were maintained in acyclovir (100µg/ml) to prevent unchromatinized lytic DNA synthesis. Shown are mean ± SD values from n=3 replicates. Blots are representative of n=3 replicates.

MR effects on epigenetic reader and writer expression could also affect the host and EBV epigenomes. Indeed, RNAseq revealed that mRNAs encoding multiple epigenetic reader and writers were highly perturbed by MR in both P3HR-1 and Rael cells (Figure 2C). MR significantly decreased levels of the initiator DNA methyl transferase DNMT3B and maintenance of DNA methylation enzymes UHRF1 and DNMT1. By contrast, MR increased mRNA levels of the TET2 and TET3 demethylases, which have important roles in support of EBV latency III gene expression and in suppression of EBV lytic genes (Guo et al., 2020a; Guo et al., 2020b; Lu et al., 2017; Wille et al., 2017b; Zhang et al., 2020) (Figure 2C). MR instead increased levels of the initiator DNA methyltransferase DNMT3A, whose overexpression alone is not sufficient to restrict LCL latency III (Guo et al., 2020b). Similarly, expression of the H3K9me2 histone methyltransferase G9a (encoded by *EHMT2*) was decreased by MR. Effects on G9a protein abundance were also evident on the protein level (Figure 2E). Since H3K9me2 has important roles in propagation of DNA methylation marks by the epigenetic enzyme UHRF1, MR-driven loss of G9a may synergistically or additively hypomethylate Burkitt cell genomes. However, histone demethylase KDM2B was significantly induced. Taken together, these results suggest that MR may alter Burkitt host and viral epigenomes both by effects on the methylation potential as well as by tilting the epigenetic reader/writer landscape towards hypomethylation.

MYC is a major regulator of B-cell immunometabolism (Stine et al., 2015). Notably, RNAseq highlighted that MYC expression was strongly suppressed by MR, which we confirmed by immunoblot analysis (Figure 2F-G). Given our recent finding that MYC represses the EBV lytic switch (Guo et al., 2020a), MR effects on MYC abundance may contribute to EBV lytic reactivation, together with hypomethylation. Pronounced MR effects on MYC and G9a expression raised the question of how MR more globally alters EBV-infected B-cell translation. Notably, diminished SAM levels can trigger SAMTOR-mediated mTORC1 repression (Gu et al., 2017). We therefore examined MR effects on protein translation levels, using a puromycin pulse labeling assay, in which puromycin incorporation into elongating nascent chains is visualized by an anti-puromycin antibody (Schmidt et al., 2009). Interestingly, whereas MR strongly reduced protein translation levels in multiple diffuse large B-cell lymphoma (DLBCL) and an acute lymphocytic leukemia cell line, reduction of media methionine concentration by 90% for 72 hours did not strongly impair translation in the three EBV+ Burkitt cell lines examined (Figure 2G). These results are consistent with the observation that MR induces expression of EBV latency and lytic genes on the protein level, and raise the question of whether EBV may alter SAMTOR sensing in Burkitt cells.

Much remains to be learned about how cross-talk between metabolic and epigenetic pathways control viral gene expression, including that of EBV. To gain insights into methionine metabolism roles in control of Burkitt EBV genomic methylation, we performed methylated DNA immunoprecipitation (MeDIP) and chromatin immunoprecipitation (ChIP) analysis of cells grown in 100 vs 10 μM methionine. As expected, the EBV *C* and *LMP* latency promoters, as well as *BZLF1* and *BMRF1* lytic promoters, were highly methylated under methionine replete conditions (Figure 2H**, S3B**). By contrast, incubation in 10 μM methionine for three days significantly decreased DNA methylation levels at each of these key EBV promotors (Figure 2H**, S3C**). Since EBV genome methylation is critical for maintenance of latency I, these results indicate that EBV+ Burkitt cells are reliant on methionine pathway flux to silence LMPs and to maintain latency.

EBV genomic histone methylation marks exert further control over latency and lytic gene expression, with H3K9me3 and H3K27me3 marks exerting important roles in silencing of lytic and latency III genes (Guo et al., 2020b; Ichikawa et al., 2018; Murata and Tsurumi, 2013). To define MR effects on these repressive chromatin marks, we performed ChIP-qPCR on nuclei from cells grown in 100 vs 10 μM methionine. MR significantly perturbed repressive H3K9me3 and H3K27me3 signals at the C, LMP, BZLF1, and BMRF1 promoters in both P3HR-1 and Mutu I cells. Interestingly, MR more strongly affected H3K9me3 than H3K27me3 levels at these key promotors (Figure 2I). Notably, levels of the H3K27 methyltransferase EZH2 were not significantly diminished by MR in multiple B-cell lines, including EBV+ BL (Figure 2G).

### The methionine cycle is necessary for maintenance of highly restricted EBV latency I

The methionine cycle produces the universal methyl donor SAM and also regenerates methionine from SAH via homocysteine (Cantoni, 1985; Dai et al., 2018; Sanderson et al., 2019) (Figure 3A). However, specific methionine cycle roles in control of the EBV epigenome have remained unstudied. To define whether flux through the methionine cycle is necessary for maintenance of EBV latency I in Burkitt cells, we used CRISPR/Cas9 editing to deplete MAT2A, the enzyme that catalyzes SAM production from methionine and ATP. We analyzed cells at an early timepoint following CRISPR editing, prior to the onset of cytotoxic effects of MAT2A inhibition. MAT2A targeting by either of two single guide RNAs (sgRNAs) de-repressed LMP1 expression to levels similar to those observed in Mutu III cells, a subclone of Mutu I with the latency III program (Figure 3B). As observed with MR, MAT2A depletion did not robustly induce latency III EBNA antigens, but did de-repress BZLF1 (Figure 3B). De-repressed LMP1 was biologically active, as evidenced by robust plasma membrane ICAM-1 induction by either MAT2A sgRNA (Figure 3C-D).

**Figure 3.**
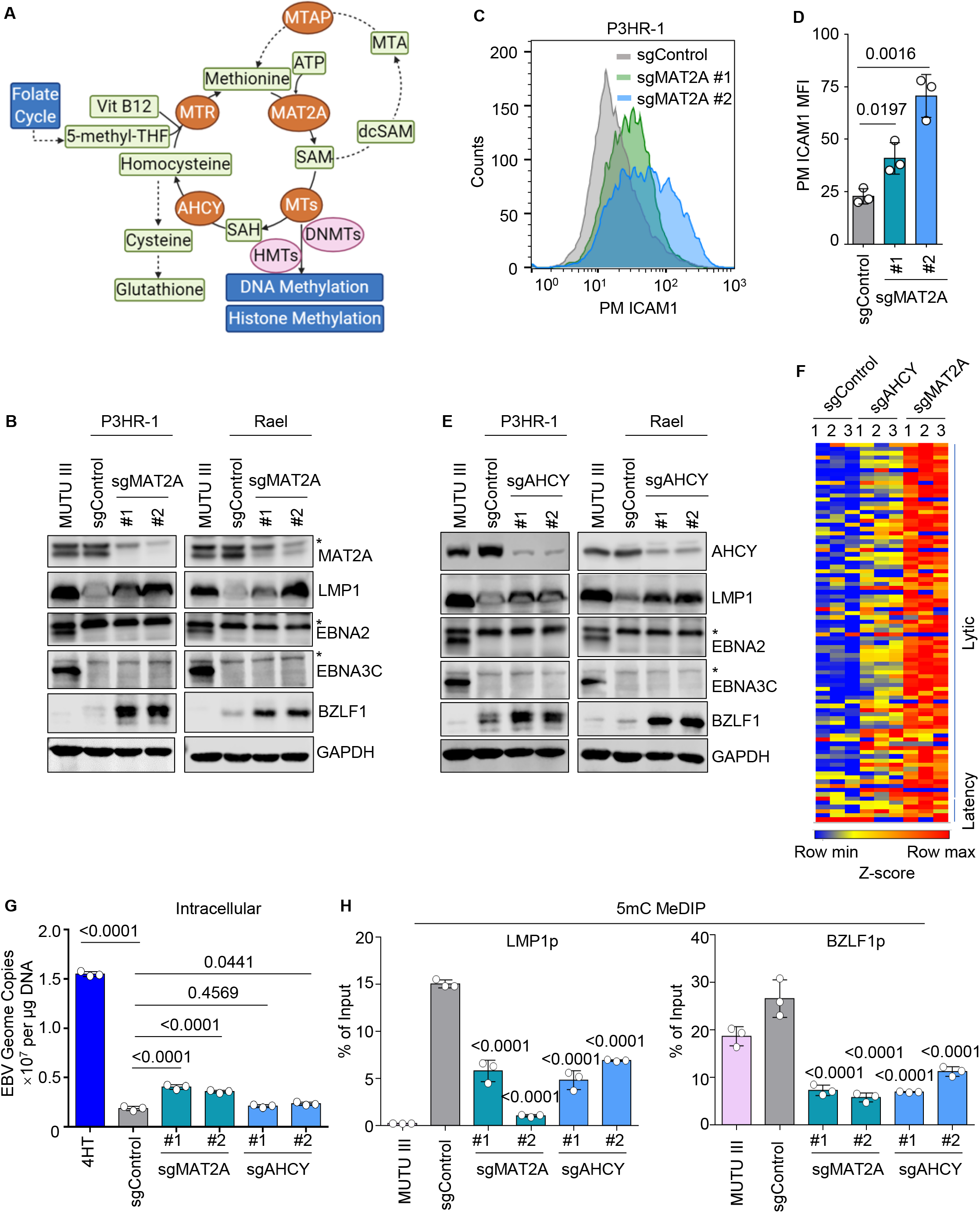
Methionine cycle flux is necessary for maintenance of EBV latency I. A) Methionine cycle schematic. MTR, methionine synthase; MAT2A, methionine adenosyltransferase 2A; MTs, methyltransferases; AHCY, adenosylhomocysteinase; dcSAM, S-adenosylmethioninamine; MTA, 5’-methylthioadenosine; MTAP, S-methyl-5’-thioadenosine phosphorylase; SAM, S-adenosylmethionine; SAH, S-adenosylhomocysteine; Vit B12, vitamin B12. Schematic figure was modified from (Sanderson et al., 2019). B) Immunoblot analysis of WCL from P3HR-1 or Rael cells expressing the indicated control or MAT2A sgRNAs. C) FACS analysis of PM ICAM1 expression in P3HR-1 cells expressing control or MAT2A sgRNAs. D) Mean ± SD PM ICAM1 MFI values from n=3 replicates of P3HR-1 cells expressing the indicated sgRNAs, as in (b). E) Immunoblot analysis of WCL from P3HR-1 or Rael cells expressing control or AHCY sgRNAs. F) RNAseq analysis of Rael cells expressing control, AHCY, or MAT2A sgRNAs. The heatmap depicts Z-score standard deviation variation from the mean value for each EBV gene across 3 replicates. G) qPCR analysis of EBV intracellular genome copy number from Rael cells expressing control, AHCY or MAT2A sgRNAs. Total genomic DNA was extracted at Day 8 post lentivirus transduction. Mean ± SD values from n=3 replicates are shown. H) 5mC MeDIP analysis of DNA from Rael cells expressing control, AHCY, or MAT2A sgRNAs, followed by qPCR using primers specific for the LMP1 or BZLF1promoters. Mean ± SEM are shown for n=3 replicates. Blots are representative of n=3 experiments.

The enzyme adenosylhomocysteinase (AHCY) converts SAH to L-homocysteine, which can be metabolized to regenerate methionine or can instead be used by the transsulfuration pathway for cysteine biosynthesis (Figure 3A). AHCY depletion strongly induced LMP1 and BZLF1 expression, suggesting that even at a step downstream of SAM generation, methionine cycle flux is critical for suppression of EBV latency and lytic antigens (Figure 3E). RNAseq identified that AHCY or MAT2A knockout de-repressed EBV latency and lytic transcripts (Figure 3F). Similar to MR, viral load analysis suggested that methionine cycle perturbation caused a largely abortive EBV lytic cycle, with comparatively small increase in EBV genome copy number (Figure 3G). Suggestive of effects at the level of the EBV epigenome, MeDIP-qPCR identified significantly loss of repressive 5mC LMP and BZLF1 promoter marks in cells depleted for MAT2A or AHCY (Figure 3H). Collectively, these results support a model in which maintenance of EBV latency I is dependent upon flow of carbon units from methionine through the methionine cycle to the EBV epigenome.

### EBV-driven methionine metabolism upregulation supports primary B cell transformation

We next asked whether an abundant methionine supply is necessary for EBV latency and lytic gene regulation in newly infected primary human B-cells. As a rationale for these studies, we noted that EBV robustly upregulates MAT2A and AHCY expression in newly B cells at the mRNA and protein levels (Figure 4A) (Wang et al., 2019a; Wang et al., 2019b). While EBV genomic methylation has key roles in control of viral latency promoters in newly infected cells and in peripheral blood B-cells of individuals with infectious mononucleosis (Ambinder et al., 1999; Guo and Gewurz, 2021; Tierney et al., 2000), EBV also highly induces the TET2 demethylase. Thus, levels of EBV genomic 5mC remain low as cells transform into LCLs (Lu et al., 2017; Wille et al., 2017a). To gain insights into how EBV alters methionine metabolism in newly infected B-cells, we performed LC/MS metabolomic profiling of B-cells at rest versus at 24 hours post-EBV infection, an early timepoint prior to the onset of newly infected B-cell proliferation. In keeping with EBV induction of methionine cycle metabolism, SAM and SAH abundances, and the cellular methylation potential, increased upon EBV infection (Figure 4B). Metabolomic pathway impact analysis identified methionine metabolism as highly rewired by EBV (Figure 4C).

**Figure 4.**
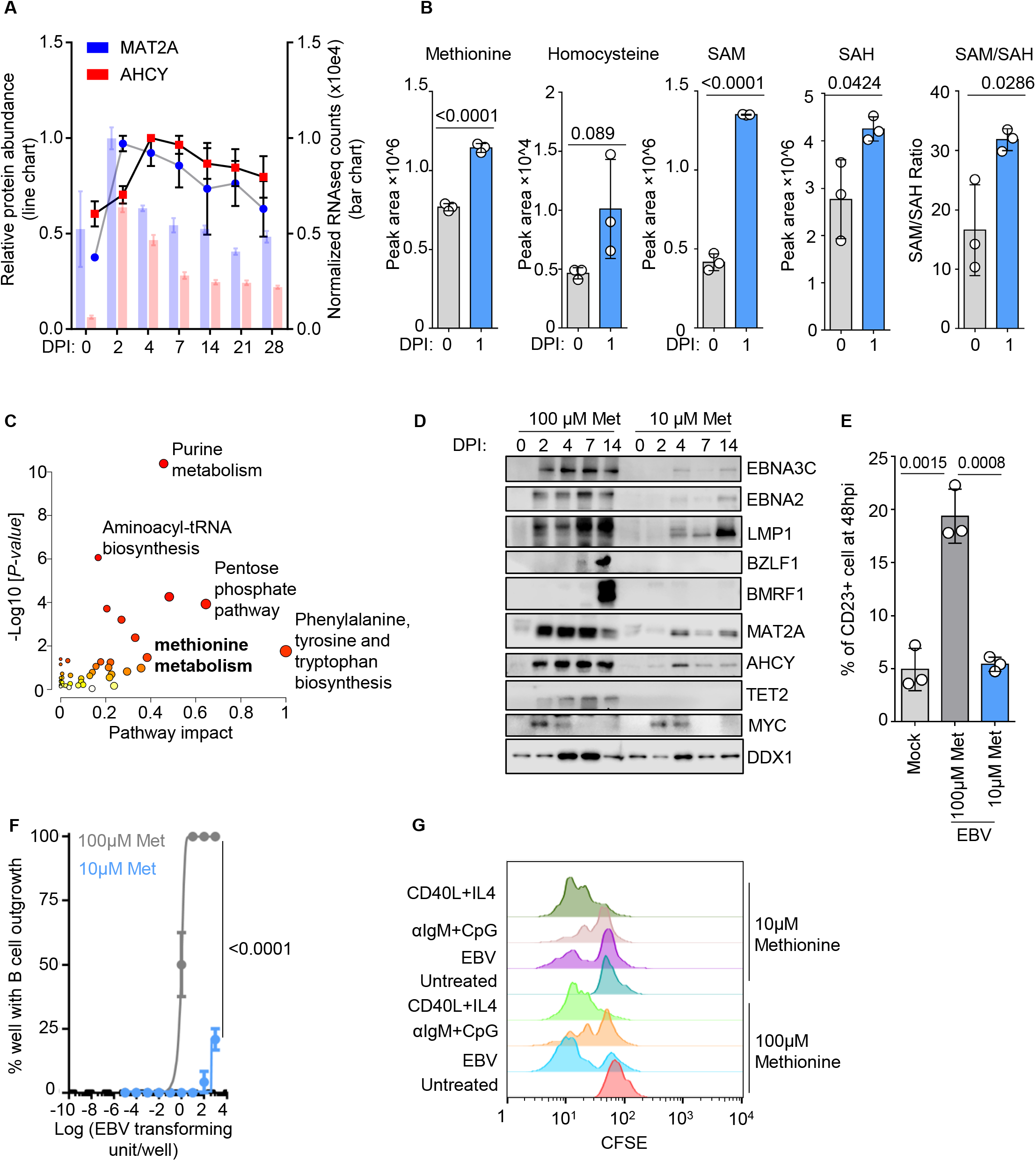
Methionine restriction inhibits EBV driven primary human B cell transformation. A) Relative mRNA (bar chart) and protein (line chart) MAT2A or AHCY levels detected by RNAseq and proteomic analysis of primary human B-cells at the indicated day post EBV infection (DPI) (Wang et al., 2019a; Wang et al., 2019b). B) Abundances of the indicated methionine cycle metabolites from primary human peripheral blood B cells at rest or 24 hours post-EBV infection. Mean ± SEM are shown for n=4 replicates. C) Metabolic pathway analysis plot showing the most strongly impacted metabolic pathways downregulated by MR in cells treated as in (A). Pathway impact value computed from MetaboAnalyst 3.0 topological analysis (x-axis) versus −Log10 (P-value) obtained from pathway enrichment analysis (y-axis). D) Immunoblot analysis of WCL from primary human B-cells at the indicated DPI and grown in 100 vs 10 μM Met, representative of n=3 replicates. E) FACS analysis of plasma membrane CD23 from EBV mock infected or EBV infected primary human B-cells at 48 hours post infection (hpi) and grown in 100 vs 10 μM Met. F) EBV transformation assays of primary human B cells grown in 100 vs 10 μM Met. Shown are fitted non-linear regression curves with means ± SEM from n=3 replicates. G) FACS CSFE dye-dilution assay of primary human B-cells untreated or stimulated as indicated by EBV MOI= 10, anti-IgM (1μg/ml), CpG (0.5mM), MEGA-CD40L (50ng/ml), IL-4 (20ng/ml) and grown in 100 vs 10 μM Met for 5 days. Representative of n=3 replicates.

We next defined MR effects on EBV latency and lytic gene expression in newly infected B-cells. Reduction of extracellular methionine concentration from 100 to 10 μM strongly perturbed EBV gene expression over the first 14 days of infection, during which EBV oncoproteins typically drive Burkitt-like and then lymphoblastoid B-cell growth (McFadden et al., 2016; Wang et al., 2019c). MR suppressed EBNA2 induction, which is essential for EBV-driven B-cell outgrowth (Pich et al., 2019), as well as EBNA2-mediated CD23 upregulation (Wang et al., 1987) (Figure 4D-E). Unexpectedly, MR also delayed EBNA2 induction to later timepoints, raising the question of how newly infected B-cells survive under MR conditions (Figure 4F). In contrast to Burkitt cells, MR failed to induce EBV lytic gene expression over the first 14 days of infection, perhaps at least in part because DNA methylation levels are considerably lower in resting and in newly infected B-cells than in Burkitt cells. Transformation assays demonstrated that MR strongly impaired LCL outgrowth (Figure 4F). These results suggest that an abundant extracellular methionine supply is necessary for epigenetic induction of the latency IIb and latency III programs and EBV-driven B-cell growth transformation into LCLs, an important model for post-transplant lymphoproliferative disease. Interestingly, MR effects on primary B-cells were also found to be stimulus specific, as MR limited outgrowth of B-cells stimulated by anti-immunoglobulin crosslinking and CpG Toll-like receptor 9 activation, but did not limit outgrowth of B-cells stimulated by CD40 ligand and interleukin-4 stimulation (Figure 4G).

### One-carbon folate metabolism is necessary for Burkitt latency I

EBV highly induces folate and one-carbon metabolism in newly infected cells and in EBV-transformed lymphoblastoid B-cells (Wang et al., 2019b), which supplies methyl groups for the methionine cycle (Ducker et al., 2017; Parsa et al., 2020; Sanderson et al., 2019) (Figure 5A). In latency I, elevated MYC levels support one-carbon metabolism in Burkitt cells. We therefore hypothesized that one-carbon folate metabolism enzymes are required for maintenance of Burkitt latency I. In support, the one-carbon enzymes MTHFD1 and MTHFD2 were hits in our recent of human genome-wide CRISPR screen for host factors that contribute to the maintenance of EBV latency I (Guo et al., 2020b) (Figure 5B).

**Figure 5.**
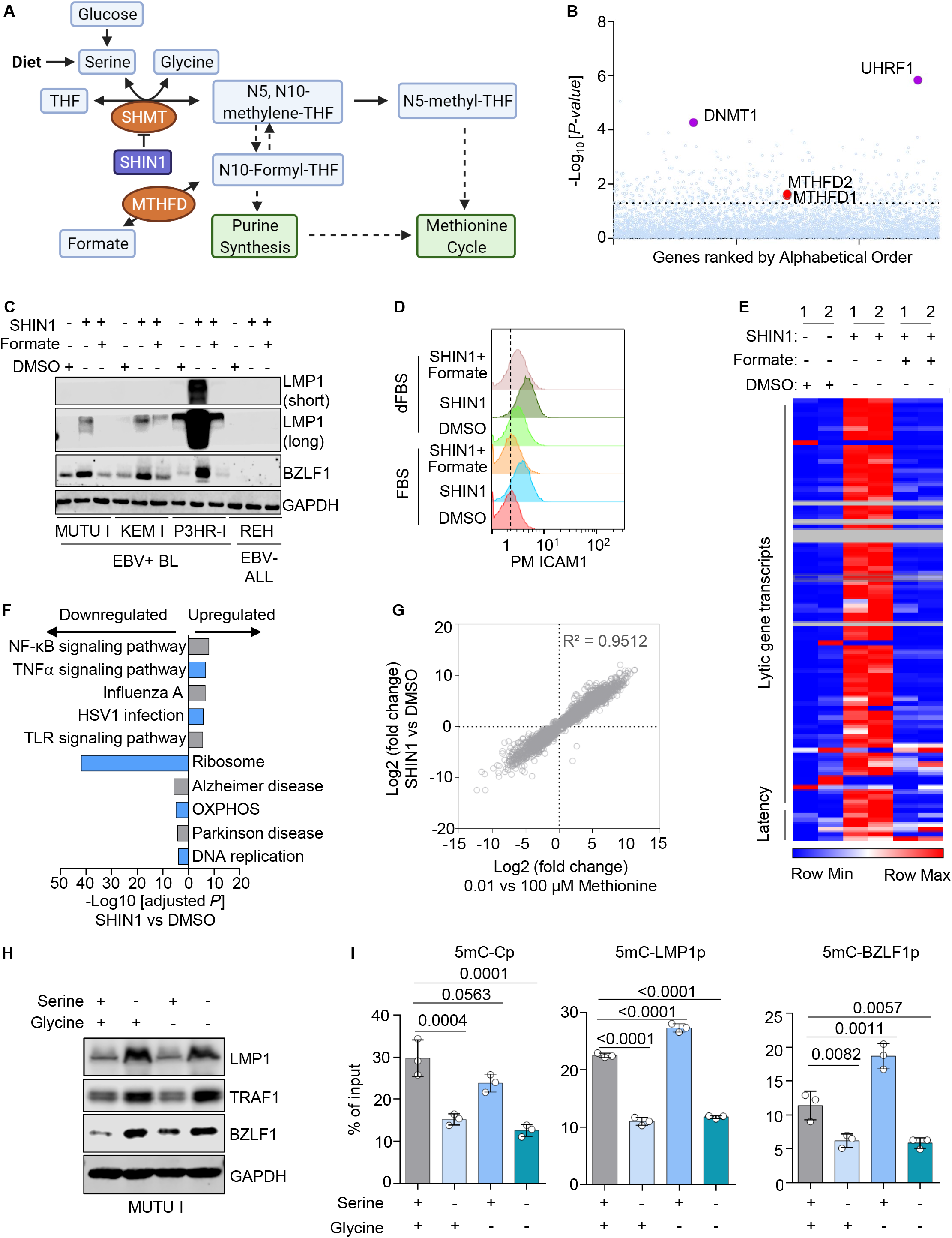
Folate cycle flux is necessary for maintenance for EBV antigens. A) Schematic of interconnected folate and methionine metabolism one carbon cycles. THF, tetrahydrofolate. SHMT, serine hydroxymethyltransferase 1 and 2. MTHFD, methylene tetrahydrdofolate dehydrogenase 1 and 2. B) Genome-wide CRISPR screen results for host factors important for maintenance of latency I(Guo et al., 2020b). Plots are representative of n = 3 biologically independent values. P-values were determined by one-sided Fisher’s exact test. Values for one carbon metabolism enzymes MTHFD1 and MTHFD2 are highlighted in red. C) Immunoblot analysis of WCL from the indicated B-cells treated with DMSO, SHIN1 (10µM), or SHIN1(10µM) and sodium formate (1mM) for 3 days. Representative of n=3 replicates. D) FACS analysis of PM ICAM1 expression in Mutu I cells treated with DMSO, SHIN1 (10µM) or SHIN1 (10µM) and sodium formate (1mM) in cells cultured with RPMI and undialyzed (bottom) versus dialyzed (top) FBS for 3 days. E) RNAseq heatmap analysis of Rael cells treated with DMSO, SHIN1 (10µM) and/or sodium formate (1mM) for 3 days, as indicated. The heatmap depicts Z-score standard deviation variation from the mean value for each EBV gene across 2 replicates. F) Enrichr pathway analysis of gene sets significantly upregulated in SHIN1 versus DMSO-treated Rael, as in (E). Enrichr adjusted p-values from n=2 samples. G) Scatter plot showing log2 fold changes in mRNA abundance in SHIN1 versus DMSO-treated Rael (y-axis) versus log2 fold changes in mRNA abundances in Rael grown in 100 vs 10 mM methionine for three days, as in Fig. 1. H) Immunoblots of WCL from Mutu I cells cultured in serine and glycine free RPMI and dialyzed FBS for four days, with add back of serine and/or glycine, as indicated. Representative of n=3 replicates. I) 5mC MeDIP analysis of Mutu I cultured in replete, serine and/or glycine depleted media for four days, as indicated, followed by qPCR using primers specific for LMP1p, BZLF1p or Cp. Mean ± SEM are shown for n=3 replicates. See also **Figure S4**.

To test whether one-carbon folate metabolism is necessary for Burkitt latency I, we blocked the cytosolic and mitochondrial pathways with the highly selective serine hydroxymethyl-transferase (SHMT) antagonist SHIN1 (Figure 5A) (Ducker et al., 2017). SHIN1 is a folate-competitive inhibitor that blocks the cytosolic SHMT1 and mitochondrial SHMT2 isoforms, and their functional redundancy may have precluded either from scoring in our CRISPR screen. SHIN1 de-repressed LMP1 and BZLF1 expression in three EBV+ Burkitt cell lines. Importantly, latency I was maintained when SHIN1 was added together with the one-carbon pathway product formate, suggesting on-target SHIN1 effects on EBV gene expression (Figure 5C). SHIN1 induced LMP1 target gene ICAM1 plasma membrane expression in Mutu I cells, an effect that was rescuable by formate. Interestingly, SHIN1 more strongly induced ICAM1 expression when cells were grown with dialyzed fetal bovine serum to reduce extracellular formate levels (Figure 5D**, S4A**). Similar results were obtained in Rael Burkitt cells (**Figure S4B-C**).

To identify SHIN1 effects on global EBV gene expression, we performed RNAseq. Reminiscent of results obtained with MR, SHIN1 induced expression of most EBV latency and lytic genes, and this effect was rescued by formate (Figure 5E). Gene ontology analysis indicated that SHIN1 significantly induced NF-κB target genes, consistent with induction of LMP1 signaling (Figure 5F). Cross-comparison of RNAseq datasets from MR and SHIN1-treated Burkitt Rael cells identified nearly identical transcriptome-wide responses (R^2^= 0.9512), suggesting that SHIN1 effects on host and viral gene expression are likely dependent on methionine cycle metabolism at this timepoint (Figure 5G).

SHMT1 and SHMT2 catalyzes the reversible conversion of serine and tetrahydrofolate to glycine and 5,10-methylene tetrahydrofolate (Figure 5A). Importantly, serine and glycine are not essential amino acids. We therefore asked whether restriction of either serine, or as a control glycine, alters EBV gene expression. Mutu I cells were cultured in serine and glycine free RPMI with dialyzed fetal calf serum. Consistent with SHIN1 effects, serine but not glycine restriction de-repressed LMP1 and BZLF1, effects that were rescued by serine add back (Figure 5H). Serine, but not glycine restriction, hypomethylated the *LMP1*, *BZLF1* and to a somewhat lesser extent *C* promoters (Figure 5I). These results are consistent with a model in which methionine and serine supply interconnecting one-carbon folate and methionine cycles that cross-talk with the EBV epigenome to maintain the Burkitt B-cell latency I state.

### Dietary MR alters the Burkitt metabolome and de-represses EBV antigens *in vivo*

Since methionine is an essential amino acid, dietary methionine restriction can be used to significantly reduce plasma methionine levels, including in murine models (Gao et al., 2019). Recent murine studies indicate that dietary MR has minimal impact on immune cell homeostasis for up to 8 weeks (Roy et al., 2020). Given our *in vitro* results, we hypothesized that Burkitt tumors are dependent upon abundant methionine supply *in vivo* to regulate the EBV epigenome and viral gene program. As a pilot study, three non-obese diabetic/severe combined immunodeficiency (NOD-SCID) NSG mice were fed either a control diet with a standard 0.86% weight by weight (w/w) methionine concentration or a MR diet with 0.086% w/w methionine, but that contained balanced total amino acid levels (**Figure S5A**). At one week, dietary MR lead to a nearly 50% reduction in plasma methionine levels (**Figure S5B),** consistent with recent mouse model MR studies (Epner et al., 2002; Roy et al., 2020). At 2 weeks after initiation of control or MR diet, Mutu I cells were implanted into mouse flanks. Body weight did not differ significantly between mice on control or MR diets over the next three weeks, suggesting that either diet was generally well tolerated (**Figure S5C**). However, tumor volumes were smaller at each timepoint in mice on the MR diet (**Figure S5D**).

We next tested MR effects on EBV+ Burkitt cell metabolomes, viral gene expression and the EBV epigenome. Mutu I tumors were implanted and allowed to grow for two weeks in twelve NSG mice fed the control diet. At that point, six mice were continued on the control diet and the other six were switched to the MR diet. At weeks four and five, mice were humanely sacrificed and samples were obtained (Figure 6A). Dietary MR significantly reduced tumor and plasma methionine and methionine sulfoxide (the oxidized form of methionine) levels at week five. Likewise, MR reduces tumor SAM and SAH levels (Figure 6B-C**, S6A**). Body weights were similar on control and MR diet fed mice (**Figure S6B**).

**Figure 6.**
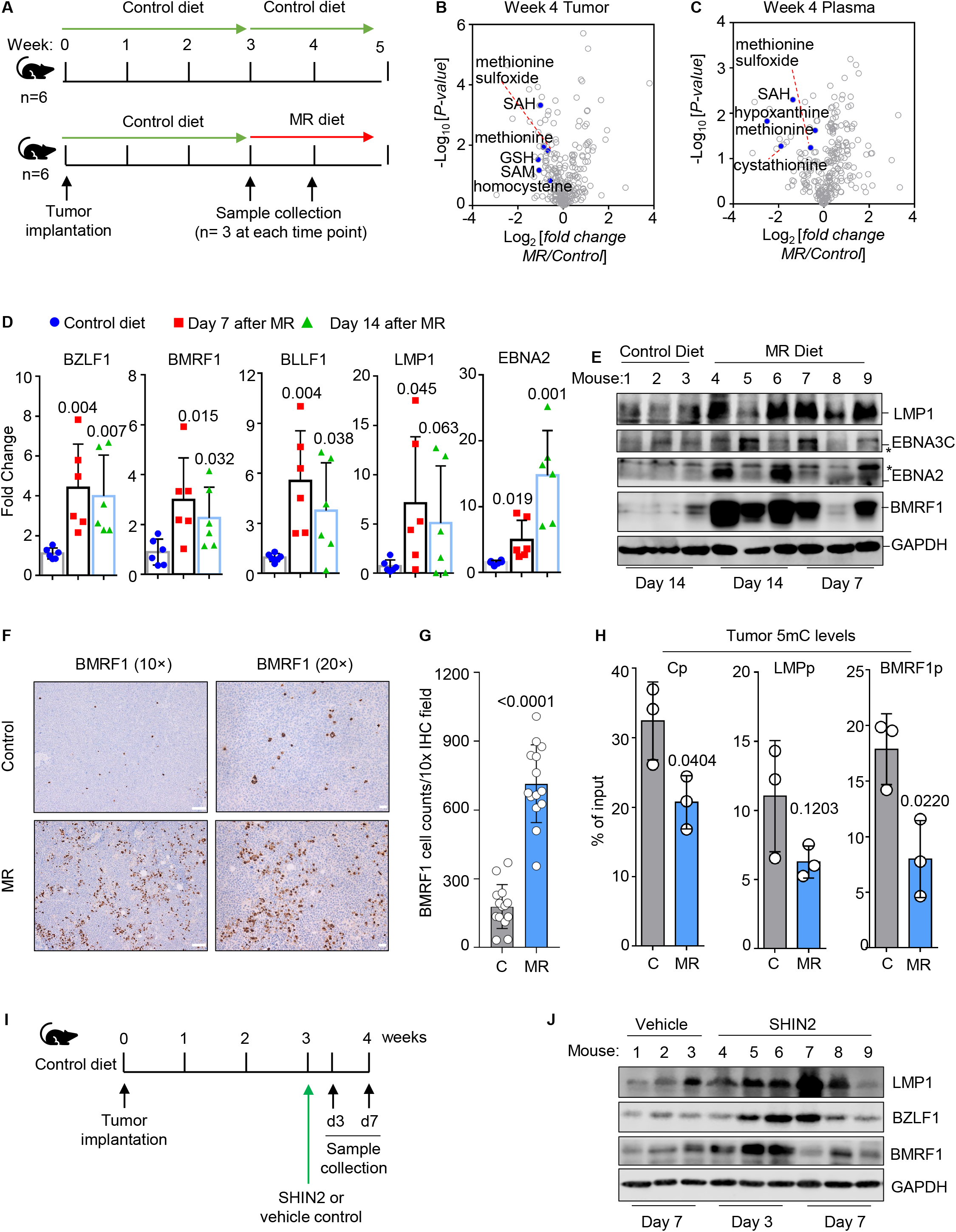
Dietary MR or pharmacological folate cycle blockade de-represses EBV immunogens *in vivo*. A) Schematic of mouse xenograft control versus MR diet intervention. B-C) Volcano plot analysis of −Log10 (p-value) versus Log2 foldchange of metabolite abundances in n=3 tumors (B) or plasma samples (C) collected from mice fed MR vs control diets for 14 days. D) qRT-PCR analysis for the indicated EBV genes from tumors explanted from mice fed control diet for 14 days or MR diets for 7 or 14 days. Mean ± SEM are shown from n=6 tumors in each group. E) Immunoblot analysis of WCL from tumors explanted from control or MR diet fed mice. F) Immunohistochemical analysis of BMRF1 expression in tumors of mice fed control or MR diet for two weeks. G) Mean ± SEM numbers of BMRF1+ cells per 10X field as in (F). H) 5mC MeDIP analysis of DNA isolated purified from tumors of mice fed control or MR diets for two weeks. Mean ± SEM values from n=3 replicates are shown. I) Schematic of Burkitt xenograft SHIN2 experiment. Mice were treated with vehicle or SHIN2 200mg/kg intraperitoneally. Samples were collected at days 3 or 7 (d3 or d7) post-treatment. J) Immunoblot analysis of WCL from tumors taken from vehicle or SHIN2-treated mice at the indicated times. P-values were determined by two-tailed Student’s t test with the two-sample unequal variance assumption. Blots in E and J are representative of n=3 replicates. See also **Figures S5, S6 and S7**.

To define dietary effects on EBV gene expression, qPCR was performed on explanted tumors from mice fed control versus MR diets. At both days 7 and 14, tumors from MR-fed mice exhibited significantly increased levels of the lytic immediate early BZFL1, early BMRF1 and interestingly also late BLLF1 genes. LMP1 mRNA levels were similarly de-repressed (Figure 6D). Immunoblots identified robust protein level upregulation of BMRF1 and LMP1 in tumors taken from five of six dietary methionine restricted mice. EBNA2 and EBNA3C were also de-repressed in a subset of tumors (Figure 6E). At a single cell level, immunohistochemical analysis demonstrated robust BMRF1 de-repression in a significant number of tumor cells (Figure 6F-G**, S6C**). MeDIP-qPCR was performed to define dietary effects on repressive 5mC DNA marks at the C, LMP and BMRF1 promoters. At each of these viral promoters normally silenced by DNA methylation in Burkitt tumors, two weeks of the MR diet significantly reduced 5mC levels, suggesting that dietary intervention alone can impact tumor methionine supply to the point of altering the EBV genomic CpG methylator phenotype (Figure 6H).

Finally, we tested effects of the SHMT1/2 inhibitor antagonist SHIN2 in the Mutu I xenograft model (Figure 6I). SHIN2 is a second generation SHIN1 derivative with improved pharmacokinetic properties (García-Cañaveras et al., 2021). Tumors were implanted and allowed to expand for three weeks in control diet fed mice. Mice were then given a single intraperitoneal injection of SHIN2 (200mg/kg) or vehicle control. Mice were then humanely sacrificed and tumor samples were analyzed at 3 and 7 days post-SHIN2 dosing. Similar to effects observed with MR, immunoblot analysis revealed LMP1, BZLF1 and BMRF1 de-repression, particularly at day 3 post-SHIN2 administration (Figure 6J**, S7A-B**). SHIN2 effects on LMP1 induction remained evident in two of three tumors at Day 7 post-dosing (Figure 6J). Taken together, these results support a model in which flux through interconnecting folate one-carbon and methionine cycle metabolism is required for maintenance of the highly restricted latency I program in Burkitt tumors.

## Discussion

Despite its position at the nexus of host cell metabolism and epigenetic reactions, little has been known about methionine cycle effects on the EBV lifecycle, or on virus/host interactions more generally. Building upon recent findings that EBV highly remodels host B-cell metabolic pathways, our results highlight key roles of interconnecting methionine and serine metabolism one carbon cycles (Maddocks et al., 2016) on the double stranded DNA EBV epigenome. Consequently, perturbation of B-cell methionine uptake or metabolism strongly altered viral gene expression and exerted major effects on the maintenance of Burkitt cell EBV latency as well as on EBV-mediated primary B-cell growth transformation (Figure 7).

**Figure 7.**
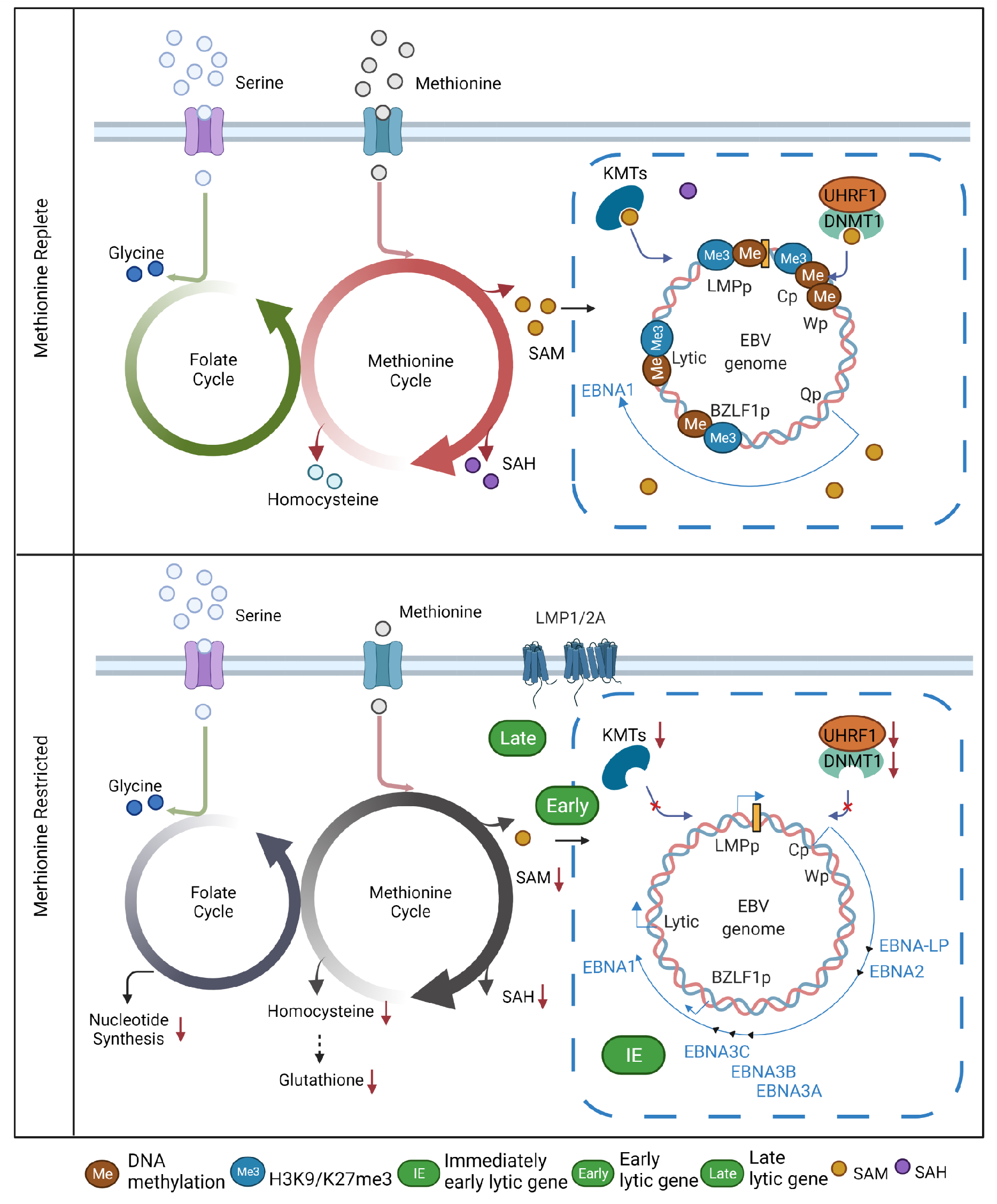
Schematic model of interconnected folate and methionine metabolism cycle roles in control of the Burkitt B-cell EBV epigenome, latent and lytic gene expression.

Burkitt lymphomas are the fastest growing human tumor (Longnecker et al., 2013; Thorley-Lawson and Allday, 2008). With each cell cycle, host factors must resynthesize and epigenetically reprogram the 10-400 EBV genomes per cell. DNA methylation has critical roles in silencing latency III and lytic antigens, which enables Burkitt tumor cells to evade antiviral immune responses (Ambinder et al., 1999; Bergbauer et al., 2010; Gewurz et al., 2021; Guo et al., 2020b). Our observations suggest that continual methionine and folate methyl cycle flux is required to meet the high Burkitt B cell demand for methyl donor groups. Superimposed effects of methionine restriction on host epigenetic enzyme expression, including down-modulation of UHRF1 and DNMT1 and upregulation of TET demethylases, likely further contributes to EBV genomic hypomethylation. It will be of interest to define whether methionine and folate metabolism have similarly important roles in other EBV disease contexts, including in nasopharyngeal and gastric carcinomas, which exhibit extreme hypermethylation and highly restricted forms of EBV latency.

MR or CRISPR methionine cycle perturbation similarly altered 5mC and histone methylation marks at the EBV *C* and *LMP* promoters. Interestingly, at the mRNA level, we observed upregulation of messages encoding EBNA3A, 3B and 3C, but not EBNA2 or LP mRNAs, even in Rael cells which have intact *EBNA2* and *EBNA-LP* genes. Since EBNA2, LP and 3 are expressed from a common *C* or *W* promoter driven transcript, we speculate that methionine metabolism also has important roles in EBNA post-transcriptional regulation, perhaps at the level of splicing. In support, methionine-derived SAM is the methyl donor for RNA methylation reactions, and RNA methylation is increasingly recognized as a key regulator of EBV-encoded mRNAs (Dai et al., 2021; Lang et al., 2019; Xia et al., 2021; Zheng et al., 2021). EBNA2 mRNAs are modified by N6-methyladenosine (m6A) marks, and knockdown of the m6A writer METTL3 decreases EBNA2 expression levels (Zheng et al., 2021). Dietary MR variably induced EBNA2 and 3C in Burkitt xenografts, albeit less robustly than LMP1 (Figure 6), suggesting that these immunogenic EBNA antigens may be more inducible by MR *in vivo,* perhaps due to additional microenvironmental effects.

It is noteworthy that MR robustly induced LMP1 and LMP2A expression, even in the absence of EBNA2. While LMP1 and LMP2A expression can be driven by the EBV lytic cycle (Yuan et al., 2006), MR strongly induced LMP1 even in CRISPR BZLF1 knockout cells, suggesting latency promoter-driven expression (**Figure S2E-F**). Yet, little is presently known about what drives LMP1 and LMP2A oncoprotein expression in the latency II program. It will therefore be of interest to determine the MR-driven host transcription factor(s) that induce Burkitt cell latent membrane protein expression. Likewise, it will be of interest to determine methionine levels in germinal centers, where EBV-infected B-cells switch to latency II and from which EBV+ Hodgkin lymphomas with latency II expression arise. Our results raise the possibility that germinal center methionine concentration may contribute to EBV latency program control.

MR robustly induced EBV immediate early and early lytic genes, but only modestly induced late genes. This observation raises the possibility that methionine metabolism supports EBV lytic DNA replication, which is required for EBV late gene expression (Chakravorty et al., 2019; Kenney and Mertz, 2014; Li et al., 2018). An abundant supply of methionine, or more likely methionine-derived SAM, may be required for modification of host or viral targets with obligatory roles in EBV lytic DNA replication, for instance to drive RNA or protein methylation reactions. It will be of interest to test whether serine metabolism is similarly important for EBV lytic DNA synthesis, such as to provide one carbon units and ATP for nucleotide synthesis (Fan et al., 2014; Huhtanen and Pensack, 1967; Maddocks et al., 2016).

Cancer cells alter methionine metabolism and increase their reliance on extracellular methionine, known as the Hoffman effect (Hoffman et al., 2019). Our findings suggest that EBV likewise subverts methionine metabolism pathways to support both the lytic and latent cycles in B-cells. Given their metabolism regulatory roles, we speculate that EBNA2 and MYC are important methionine metabolism regulators in newly infected cells. In support, EBNA2 is the only EBV protein coding gene required for primary B-cell growth transformation over the first 8 days of infection (Pich et al., 2019). Aberrantly elevated MYC levels may similarly drive Hoffman metabolism in Burkitt cells with the latency I program.

Epigenetic enzymes are regulated by substrate and product concentrations (Dai et al., 2020; Lu and Thompson, 2012). Dietary methionine restriction achieved approximately 50% reduction in plasma methionine, significantly reduced xenograft methionine abundance and tumor volumes, and diminished EBV genomic 5mC levels at the *C*, *LMP1* and *BMRF1* promoters (Figure 6**, S5-6**). Yet, MR diets for up to eight weeks have minimal impact on spleen or lymph node T, B or NK cell numbers, despite similar reduction in plasma methionine to what we observed (Epner et al., 2002; Roy et al., 2020). MR also did not significantly affect CD4+ T-helper 1 (Th1) lymphocyte proliferation or histone epigenetic activation marks. This may relate at least in part to the observation that antigen activated T-cells upregulate plasma membrane methionine transporter expression (Sinclair et al., 2019). These observations raise the possibility that EBV latently infected B-cells are particularly sensitive to MR, and that a therapeutic window may exist for implementing MR together with anti-EBV T-cell immunotherapies.

Recent epidemiological and immunological studies suggest that EBV is a major trigger for multiple sclerosis (MS), in which autoreactive immune responses attack nerve fiber myelin sheaths (Bjornevik et al., 2022; Lanz et al., 2022; Robinson and Steinman, 2022). Interestingly, dietary methionine restriction ameliorates neuroinflammation and disease onset in the mouse experimental autoimmune encephalomyelitis (EAE) model of MS. 33% of mice on an MR diet exhibited symptoms at two weeks post EAE induction challenge, as compared with 90% in animals fed a control diet (Roy et al., 2020). Mechanistically, dietary MR reduces inflammatory T helper 17 (Th17) cell H3K4me3 activating promoter marks, proliferation and cytokine production (Roy et al., 2020). By contrast, administration of LCLs, or even an EBNA1 peptide implicated in molecular mimicry with the adhesion molecule GlialCAM, exacerbates EAE (Lanz et al., 2022; Polepole et al., 2021). It will therefore be of considerable interest to investigate dietary MR effects on EAE in humanized mouse models of EBV infection. Use of humanized mouse models carrying MS genetic risk factor HLA-DRB1*15:01 (HLA-DR15) allele may be particularly informative (Zdimerova et al., 2021). Given our observation that MR impairs the outgrowth of newly EBV infected B-cells, it is plausible that MR could synergistically reduce B-cell EBNA1 abundance as well as pathologic Th17 responses.

Current approaches to Burkitt lymphomas therapy do not harness the presence of viral genomes in EBV+ tumors. Consequently, use of high intensity chemotherapy causes substantial side-effects and has limited efficacy. While there is growing interest in use of immunotherapy approaches to treat B-cell lymphomas, the majority of EBV+ Burkitt tumors evade immune detection by restricting expression of all but EBV-encoded protein antigen. Our results raise the possibility that lowering extracellular methionine levels through dietary MR, or perhaps lowering methionine levels by orally-administered methioninase (Yamamoto et al., 2022), may sensitize tumor cells to attack by anti-EBV cytotoxic T-cells. Indeed, B-cell LMP1 expression can elicit potent antitumor T-cell responses (Choi et al., 2021). Similarly, MR mediated de-repression of EBV lytic antigens has the potential to sensitize tumor cells to highly cytotoxic effects of ganciclovir, an antiviral agent that is activated by an EBV-encoded lytic cycle early gene kinase (Meng et al., 2010). Given our observation that MR restrains EBV-driven B-cell immortalization, it will also be of interest to investigate whether dietary MR can be used in prophylaxis approaches for the prevention of EBV+ post-transplant lymphoproliferative diseases, particularly in high risk acute post-transplant settings.

In summary, our results suggest that EBV-infected B-cells are highly sensitive to perturbation of extracellular methionine or serine concentrations, or blockade of interconnecting methionine and folate one-carbon metabolism cycles. Disruption of the flow of methyl groups from methionine onto the EBV epigenome de-represses the Burkitt B-cell highly restricted latency I state. MR highly induced EBV latent membrane proteins and an abortive lytic cycle, largely comprised of highly immunogenic (Taylor et al., 2015) immediate early and early antigens. Conversely, methionine restriction significantly impairs EBV-mediated primary B-cell growth transformation, suggesting that methionine metabolism also exerts key effects on EBV latency oncoprotein expression even in the absence of CpG island methylator phenotypes.

## ACKNOWLEDGEMENTS

We thank Jaap Middeldorp for generously sharing the anti-BMRF1 antibody OT14E and the anti-LMP1 antibody OT21C. We thank Adam Friedman and Raze therapeutics for supplying SHIN1. We thank John Asara and the Beth Israel Mass Spectrometry Core Facility. We thank the Cornell University College of Veterinary Medicine PATh core facility. This work was supported by NIH R01 AI137337, CA228700 and AI164709, a Burroughs Wellcome Career Award in Medical Sciences to BEG, and a Lymphoma Foundation Fellowship to RG. All RNAseq datasets will be published in the GEO omnibus upon manuscript publication, GSE190937 and GSE190975.

## AUTHOR CONTRIBUTIONS

R.G., J.H.L., Y.C.Z., M.L., Z.X.L., Y.W., V.T.A., R.P. performed the experiments. R.G., L.G.R and B.G. designed the experiments. R.G. performed bioinformatic analysis. R.G. and B.E.G. wrote the first draft. All authors analyzed the results, read and approved the manuscript.

## RESOURCE AVAILABLITIY

Lead Contact Further information and requests for resources and reagents should be directed to and will be fulfilled by the Lead Contact, Rui Guo, Benjamin Gewurz.

### Materials Availability

All plasmids and cell lines generated in this study will be made available on request.

### Data and Code Availability

All RNA-seq datasets have been deposited to the NIH GEO omnibus. The accession number for the RNA-seq dataset reported in this paper is database: GSE190937 and GSE190975. Figures were drawn with commercially available GraphPad, Biorender, Microsoft Powerpoint.

## EXPERIMENTAL MODEL AND SUBJECT DETAILS

### Cell lines and reagents

The Burkitt cell lines Mutu I and KEM I were obtained from Dr. Jeffrey Sample. Burkitt cell lines P3HR-1 clone 16, EBV+ Akata, EBV-Akata and Daudi BL were obtained from Elliott Kieff. Rael BL cells, HBL1, TMD8 DLBCL cells were obtained from Ethel Cesarman. The B-ALL REH cell line and Farage EBV+ DLBCL were obtained from ATCC. Cell lines with stable *Streptococcus pyogenes* Cas9 expression were generated by lentiviral transduction and blasticidin selection (5 µg/ml), as previously described(Ma et al., 2017a). Unless otherwise indicated, all cell lines used in the study were Cas9+. Cells were cultured in a humidified incubator at 37°C with 5% CO2 and routinely tested and certified as mycoplasma-free using the MycoAlert kit (Lonza). B cells were grown in RPMI 1640 medium (Gibco, Life Technologies) with 10% fetal bovine serum (FBS, Gibco) or with dialyzed FBS where indicated (Gibco). 293T were obtained from ATCC and grown in Dulbecco’s Modified Eagle’s Medium (DMEM) with 10% FBS. For selection of transduced cells, puromycin was added at the concentration of 3 µg/ml. The SHMT1/2 dual inhibitor SHIN1 was obtained from Raze Therapeutics and used at 10µM. Sodium formate (Sigma-Aldrich) was used at 1M. Acyclovir was used at 100 µg/ml. All cell lines were routinely tested for mycoplasma by the Lonza Mycoalert kit, according to the manufacturer’s instructions. Antibodies used in the study was listed in Key Resources table.

### *In vitro* methionine restriction

B-cells cultured in RPMI-1640 with 10% regular FBS were washed with PBS three times and then resuspended in methionine free RPMI-1640 (ThermoFisher Cat # A1451701) supplemented with 10% dialyzed FBS (ThermoFisher Cat #26400044). L-methionine was added to the cell culture to the indicated concentrations and cells were grown for the indicated times, typically 72 hours.

### *In vitro* serine and/or glycine depletion

Mutu I cells cultured in replete media (RPMI-1640 with 10% regular FBS) were washed with PBS three times and then resuspended in serine, glycine and glucose free RPMI-1640 (Teknova Cat# R9660-02), supplemented with 11.1mM D-glucose and 10% dialyzed FBS. For serine depletion, 133µM glycine was added. For glycine depletion, 286 µM L-serine was added. For dual serine/glycine depletion, vehicle control was added. For replete media conditions, 286 µM L-serine and 133µM glycine was added.

### CRISPR/Cas9 editing

CRISPR/Cas9 engineering was performed in cells with stably Cas9 expression, using Broad Institute Brunello library sgRNA sequences. sgRNA oligos were obtained from Integrated DNA Technologies and cloned into the pLentiGuide-Puro vector (Addgene plasmid #52963, a gift from Feng Zhang). Lentiviruses were produced in 293 cells by co-transfection of pLentiGuid-puro with psPAX2 and VSV-G packaging. At 24 hours post transfection, cell culture media was changed to RPMI-1640+10% FBS. Two rounds of lentiviral transduction were performed at 48 and 72 hours post plasmids transfection. Cells were selected by puromycin (3 µg/ml), added 48 hr post-transduction. Depletion of target gene encoded protein expression was confirmed by immunoblot.

### Immunoblot analysis

Immunoblot was performed as previously described (Ma et al., 2017b). In brief, whole cell lysates (WCL) prepared by boiling cells in 1× Laemmli buffer were separated by SDS-PAGE electrophoresis, transferred onto the nitrocellulose membranes, blocked with 5% milk in TBST buffer and then probed with relevant primary antibodies at 4 °C overnight, followed by secondary antibody incubation for 1 h at room temperature. Blots were then developed by incubation with ECL chemiluminescence for 1 min (Millipore) and images were captured by Licor Fc platform. Bands intensities were measured where indicated by Image Studio Lite Version 5.2. All antibodies used in this study were listed in the Key Resources Table.

#### Puromycin analysis of protein translation

Two million cells were seeded at 0.3 million per ml in RPMI-1640+10% FBS. Puromycin (10 μg/ml) was added for 20 min at 37°C. WCLs were prepared and analyzed by immunoblot, using an anti-puromycin monoclonal antibody to visualize newly synthesized polypeptides.

### Flow cytometry analysis

Flow cytometry was performed on a BD FacsCalibur instrument. For live cell antibody staining, 1× 10^6^ cells were washed twice with FACS buffer (PBS, 1mM EDTA, and 0.5% BSA) and then stained with fluorophore conjugated anti-gp350 or anti-ICAM1 primary antibodies for 30 minutes on ice. Labeled cells were then washed three times with FACS buffer prior to the flow cytometry. For CFSE staining, BL, REH, or primary B cells were stained with 10 µM CFSE for 15 minutes at 37°C, washed, resuspended at 100 000 cells/mL and then treated with indicated conditions. For cell cycle analysis, cells collected at day 3 post methionine restriction were fixed in 70% ethanol overnight, washed once with 1×PBS and resuspended in staining buffer (propidium iodide, 5µg/ml; RNase A, 40µg/ml; 0.1% Triton X-100 in PBS) for 30 minutes at room temperature. Cells were then analyzed by FACS. FACS data were analyzed with FlowJo V10.

### Dot blot

1µg of DNA harvested by DNeasy Blood& Tissue Kit from cells grown under indicated condition was hybridized on the nitrocellulose membrane and blotted with anti-5 methyl-cytosine monoclonal antibody or stained with 1µg/ml ethidium bromide in PBS for 10min at room temperature. The membrane was then washed with PBS three time. Dot blots were imaged on a Li-Cor Fc system, and dot intensity was quantitated by Image Studio Lite Version 5.2.

### Chromatin immunoprecipitation (ChIP)

2×10^7^ of cells were fixed with 10 ml of 1% formaldehyde in RPMI-1640+10% FBS for 10min at room temperature. The cross-linking reaction was then quenched by adding 1.425ml of 1M glycine for 10min at room temperature. Cells were washed with ice-cold PBS for three times. Then, cells were incubated with 2ml of lysis buffer (50 mM Tris pH8.1, 10 mM EDTA, 1% and 1% SDS, protease inhibitor cocktail) for 20 minutes on ice. Chromatin were fragmented with Bioruptor (Diagenode, USA) with 30s on/ 30s off condition for 30 cycles. Fragmented chromatin was then diluted with dilution buffer (16.7 mM Tris pH8.1, 1.2 mM EDTA, 167mM NaCl, 1.1% Triton-X100, 0.01% SDS, and protease inhibitor cocktail) and incubated with 5μg of the ChIP grade antibody of interest (see Key Resource Table) at 4°C overnight. Immunocomplexes were precipitated by addition of 100µl of prewashed protein A or G magnetic beads for 1h at 4°C. Beads were isolated using a magnet and washed 2 times with 10ml low salt buffer (20 mM Tris pH8.1, 2 mM EDTA, 150mM NaCl, 1 % Triton-X100, 0.1% SDS), 2 times with 10 ml high salt buffer(20 mM Tris pH8.1, 2 mM EDTA, 500mM NaCl, 1 % Triton-X100, 0.1% SDS), once with 10 ml lithium chloride buffer(10 mM Tris pH8.1, 1 mM EDTA, 0.25M LiCl, 1 % NP40, 1% deoxycholic acid, and once with 10ml TE buffer. DNA was eluted using freshly prepared elution buffer (1%SDS, 100Mm NaHCO_3_ in H_2_O). 250µl (per 100µl beads) of elution buffer was added and incubated at room temperature for 10min. The eluted DNA was then reverse crosslinked by treating with 20U protease K at 65°C overnight. After reverse crosslinking, DNA was purified by the QIAquick PCR purification kit (Qiagen) according to the protocol. ChIP assay purified DNA was quantitated by qPCR, and values were normalized to the percentage of input DNA. Primers for qPCR are listed in Supplementary Table S7.

### 5-methyl Cytosine DNA immunoprecipitation (MeDIP) and qPCR

Genomic DNA was purified using the Blood and Cell Culture DNA Mini Kit (Qiagen) and then used for MeDIP analysis, using the MagMeDIP kit (Diagenode cat # C02010021). qPCR assays were then performed as described above. Primers for qPCR are listed in Supplementary Table S7.

### Quantitative real time (qRT)-PCR

Total RNA was harvested from xenograft tumors by the RNeasy Mini Kit (Qiagen) according to the manufacturer’s instructions. Genomic DNA was removed by RNase-Free DNase Set (Qiagen) and qRT-PCR was then performed using the Power SYBR Green RNA-to-CT 1-Step Kit (Applied Biosystems) on a CFX96 Touch™ Real-Time PCR Detection System (Bio-Rad). Data was normalized to internal control 18S rRNA. Relative expression was calculated using 2-ΔΔCt method. All samples were run in technical triplicates and at least three independent experiments were performed. Primer sequences are listed in Supplementary Table S7. Primers used for detection of BZLF1, BMRF1, BLLF1, LMP1 or 18S rRNA were described previously(Lu et al., 2017).

### RNAseq analysis

Total RNA was isolated by the RNeasy Mini kit (Qiagen), following the manufacturer’s manual. An in-column DNA digestion step was included to remove the residual genomic DNA contamination. To construct indexed libraries, 1 µg of total RNA was used for polyA mRNA-selection, using the NEBNext Poly(A) mRNA Magnetic Isolation Module (New England Biolabs), followed by library construction via the NEBNext Ultra RNA Library Prep Kit (New England Biolabs). Each experimental treatment was performed in triplicate. Libraries were multi-indexed, pooled and sequenced on an Illumina NextSeq 500 sequencer using single-end 75 bp reads (Illunima) at the Dana Farber Molecular Biology core. Adaptor-trimmed Illumina reads for each individual library were mapped back to the human GRCh37.83 transcriptome assembly using STAR2.5.2b (Dobin et al., 2013). Feature Counts was used to estimate the number of reads mapped to each contig (Liao et al., 2014). Only transcripts with at least 5 cumulative mapping counts were used in this analysis. DESeq2 was used to evaluate differential expression (DE) (Love et al., 2014). DESeq2 uses a negative binomial distribution to account for overdispersion in transcriptome datasets. It uses a conservative analysis that relies on a heuristic approach. Each DE analysis used pairwise comparison between the experimental and control groups. Differentially expressed genes were identified and a p values < 0.05 and absolute fold change > 2 cutoff was used. Differentially expressed genes were subjected to Enrichr analysis which was employed to perform gene list-based gene set enrichment analysis on the selected gene subset. The algorithm used to calculate combined scores was described previously (Chen et al., 2013). P value and log2 fold change were generated with DESeq2 under default settings with Wald test and normal shrinkage, respectively. Top 5 Enrichr terms that passed the adjusted p-value cutoff were visualized using Graphpad Prism 7.

Volcano plots were built with Graphpad Prism7. Heatmaps were generated by feeding Z-score values of selected EBV genes from DESeq2 into Morpheus software (https://software.broadinstitute.org/morpheus/).

### Confocal microscopy

Cells were seeded on glass slides in PBS, air dried and then fixed with 4% paraformaldehyde (PFA) in PBS for 10 minutes. PFA was removed and fixed cells were permeabilized with 0.1% Triton-X in PBS. Slides were blocked with 1% IgG-free BSA (Sigma-Aldrich, Cat# A2058) in PBS for 30 minutes at room temperature. Cells were incubated with primary antibodies against LMP1 (monoclonal antibody OT21C, a gift from Dr. Jaap Middeldorp, 1:1000), LMP2A (ab59028, Abcam, 1:200) or EBNA2 (PE2, a gift from Dr. Elliot Kieff) in PBS containing 1% BSA for 1 hour at 37℃. Slides were then washed three times and then incubated with secondary antibodies (Alexa Fluor 488-conjugated goat anti-mouse and/or Alex Fluor 594-conjugated goat anti-rabbit, diluted 1:250 in PBS) for 1 hour at 37℃. Slides were washed three times in PBS and incubated with 100 uL of Hoechst 33258 (10 µg/mL in PBS) for 10 minutes. Cells were then washed three times with PBS. ProLong Gold anti-fade was applied to the slide, which was then sealed with a No. 1.5 coverslip. Image acquisition was performed with the Zeiss LSM 800 instrument. Image analysis was performed with the Zeiss ZEN Blue software.

### Quantification of intracellular EBV genome copy number

To measure EBV genome copy number, intracellular viral DNA was quantitated by qPCR analysis. For intracellular viral DNA quantitation, total DNA from 2×10^6^ Burkitt cells was extracted by the Blood & Cell culture DNA mini kit (Qiagen #13362). Extracted DNA was diluted to 10 ng/ul and analyzed by qPCR targeting the EBV BALF5 gene. Standard curves were made by serial dilution of a pHAGE-BALF5 miniprep DNA at 25 ng/uL. Viral DNA copy number was calculated by inputting sample Ct values into the regression equation dictated by the standard curve.

### EBV production and concentration

EBV+ Akata cells were resuspended in FBS-free RPMI 1640 at a concentration of 2–3×10^6^ cells per ml and then inducted with 10µg/ml goat anti-human IgG (Dako, Cat# A042402-2) for 72 h at 37°C. Cell culture supernatant was collected and passed through a 0.45 µm filter and then pelleted by centrifugation at 50,000 g for 2.5 hours, and then resuspended in PBS with 2% dialyzed FBS. The virus was aliquoted, stored at −80°C and thawed immediately before infection. Titer was determined experimentally by transformation assay as described below.

### Primary Human B-cells

Discarded, de-identified leukocyte fractions left over from platelet donations were obtained from the Brigham and Women’s Hospital Blood Bank. Blood cells were collected from platelet donors following institutional guidelines. Since these were de-identified samples, the gender was unknown. Our studies on primary human blood cells were approved by the Brigham & Women’s Hospital Institutional Review Board. Primary human B cells were isolated by negative selection using RosetteSep Human B Cell Enrichment and EasySep Human B cell enrichment kits (Stem Cell Technologies), according to the manufacturers’ protocols. B cell purity was confirmed by plasma membrane CD19 positivity through FACS. Cells were then cultured with RPMI 1640 with 10% FCS. Freshly isolated primary B-cells were seeded in control or MR media at the density of 1 million/ml.

### EBV Transformation Assay

Freshly isolated primary human B-cells, purified as outlined above by negative selection, were seeded into 96-well plates at a density of 500,000 cells/mL in 100 μL per well of control or MR media. Akata cell supernatant was diluted ten-fold to give a five-point dilution series. To each well, 100 μL of virus supernatant was added. Control or MR media were refreshed every 3 to 4 days by carefully aspirating 100 μL of spent media and replenishing with fresh media. At four weeks post-infection, the proportion of wells with B-cell outgrowth was plotted against the dilution of virus supernatant used per well, as previously described(Henderson et al., 1977). One transforming unit per well was defined as the amount of virus required to attain B-cell outgrowth in 62.5% of wells.

### Intracellular metabolite profiling

For P3HR-1 profiling, 3.5×10^6^ cells were seeded into a T25 flask with 10 mL of fresh control or MR media, supplemented with 10% dialyzed FBS. Three days after seeding, 3×10^6^ P3HR-1 cells were pelleted and resuspended in fresh control or MR media for 2 hours prior to intracellular metabolite extraction. For profiling of resting versus newly infected B-cells, 5×10^6^ freshly isolated primary B cells were mock infected or infected by EBV at a MOI of 10 for 22 hours. 3×10^6^ mock or EBV infected cells were pelleted and resuspended in fresh RPMI-1640 media supplemented with 10% dialyzed FBS for 2 hours prior to intracellular metabolite extraction. Cells were pelleted and washed with 5 mL of room temperature PBS. Pellets were resuspended in 1 mL of dry ice cold 80% methanol, incubated at −80°C for 30 minutes and centrifuged at 21,000 x g for 5 minutes to precipitate proteins. For mice xenograft tumors, 100mg of tumor tissues were washed once with PBC and submerged in 500 µl of 80% (vol/vol) HPLC-grade methanol (cooled to −80 °C). The tissues were smashed/grinded for 1–2 min with small pestle/tissue grinder on dry ice in the tube, vortex for 1 min at 4–8 °C and incubate for 4 hours at −80 °C. The tissues were then centrifuged at 21,000g for 10 min using a refrigerated centrifuge (4–8 °C). The supernatant was collected in pre-chilled tubes and stored at −80°C. On the day of analysis, supernatants were incubated on ice for 20 minutes and clarified by centrifugation at 21,000 x g at 4°C. At the Beth Israel Mass Spectrometry core, supernatants were dried down in a speed vacuum concentrator (Savant SPD 1010, Thermofisher Scientific) and re-suspended in 100µL of 60/40 acetonitrile/water. The samples were then vortexed, sonicated in ice-cold water for 1 minute, and incubated on ice for 20 minutes. Supernatants were collected in an autosampler vial after centrifugation at 21,000 x g for 20 minutes at 4°C. Pooled QC samples were generated by combining ∼15µL of each sample. Metabolite profiling was performed using Dionex Ultimate 3000 UHPLC system coupled to Q-Exactive plus orbitrap mass spectrometer (ThermoFisher Scientific, Waltham, MA) with an Ion Max source and HESI II probe operating in switch polarity mode. A zwitterionic Sequent zic philic column (150 x 2.1mm, 5µm polymer, part # 150460, MilliporeSigma, Burlington, MA) was used for polar metabolite separation. Mobile phase A (MPA) was 20mM ammonium carbonate in water, pH9.6 (adjusted with ammonium hydroxide) and MPB was acetonitrile. The column was held at 27°C, injection volume 5µL, autosampler temperature 4°C and LC conditions at flow rate of 0.15 mL/min were: 0min: 80% B, 0.5min: 80% B, 20.5min: 20% B, 21.3min: 20%B, 21.5min: 80% B with 7.5min of column equilibration time. MS parameters were: sheath gas flow 30, aux gas flow 7, sweep gas flow 2, spray voltage 2.80kV for negative & 3.80kV for positive, capillary temperature 310°C, S-lens RF level 50 and aux gas heater temp 370°C. Data acquisition was done using Xcalibur 4.1 (ThermoFisher Scientific) and performed in full scan mode with a range of 70-1000m/z, resolution 70,000, AGC target 1e6 and maximum injection time of 80ms. Data analysis was performed in Compound Discoverer 3.1 and Tracefinder 4.1. Samples were injected in randomized order and pooled QC samples were injected regularly throughout the analytical batch. Metabolite annotation was done base on accurate mass (±5ppm) and matching retention time (±0.5min) as well as MS/MS fragmentation pattern from the pooled QC samples against in-house retention time +MSMS library of reference chemical standards. Metabolites with CV<30% in pooled QC were used for the statistical analysis. The quality of integration for each metabolite peak was reviewed. Metabolites with p-values < 0.05, log_2_(fold change)>1 or <-1 were used for pathway analysis using MetaboAnalyst 5.0 (https://www.metaboanalyst.ca/MetaboAnalyst/ModuleView.xhtml).

### Mouse xenograft experiments

Mouse xenograft experiments were done in accordance with the Institutional Animal Care & Use Committee (IACUC# 2017-0035) of Weill Cornell Medical Center(WCMC).

Trial 1: Six NOD.Cg-Prkdcscid Il2rgtm1Wjl/SzJ (NSG) immunocompromised mice were divided into two groups, which were fed the control amino acid diet (TD01084, Envigo) or a 90% reduced methionine diet (TD200744, Envigo) for two weeks prior to bilateral xenotransplantation with a finely minced 2-3mm3 piece of EBV-related Burkitt Lymphoma, Mutu I. Mice were raised in control or MR diet for additional 3 weeks. Bilateral tumors were collected and processed 1/3 flash frozen in liquid nitrogen, 1/3 fixed in 10% formalin, and 1/3 stored in RNAlater. A tail vein pre-bleed was performed on all mice and weekly tail bleeds were performed each week from tumor implantation.

Trial 2: Twenty four NOD.Cg-Prkdcscid Il2rgtm1Wjl/SzJ (NSG) immunocompromised mice were xenotransplanted bilaterally with a finely minced 2-3mm3 piece of EBV-related Burkitt Lymphoma, Mutu-1. Twelve mice were placed on a control diet (TD200744, Envigo) for three weeks until tumor burden reached sufficient size at which point animals were randomly placed into groups and six animals were switched to a methionine restricted diet (TD200744, Envigo). Half the animals were euthanized (n=3 control diet, n=3 methionine restricted diet) at day 7 post diet change and the other half at day 14 post diet change.

The remaining 12 mice were placed on in house irradiated mouse diet (Teklad, 7912). Once tumor burden reached sufficient size 12 mice were randomized into vehicle and treatment groups (n=6/group). Either a single injection of SHIN2 (stock concentration 20mg/ml, dosing concentration 200mg/kg) or vehicle control was given intraperitoneally (IP). Half the animals were euthanized (n=3 SHIN2 Tx, n=3 vehicle control) at day 3 post injection and the other half at day 7 post injection.

Bilateral tumors were collected and processed 1/3 flash frozen in liquid nitrogen, 1/3 fixed in 10% formalin, and 1/3 stored in RNAlater. A tail vein pre-bleed was performed on all mice and weekly tail bleeds were performed each week from tumor implantation. A final cardiac bleed was performed at sacrifice. Whole blood was spun down for serum collection and frozen at −20C and then moved to −80C.

Digital caliper measurements and body weight were measured under 2.5% isoflurane anesthesia. Two-dimensional length (L) and width (W) measurements were used to extrapolate 3D volume using the formula (L x W^2^)/2. All animals were euthanized by CO2 asphyxiation at a flow rate 3.5L/min. Diet composition is listed in supplementary table S7.

### Immunohistochemistry

Tumor samples were fixed in 10% neutral buffered formalin (Sigma-Aldrich) at room temperature. After trimming, tissues in the cassettes were processed using a Tissue-Tek VIP automatic tissue processor (Sakura, Finland) with the routine overnight run and embedded into paraffin wax (Tissue-Tek). Embedded blocks were placed on the ice for a short time, and cut into 4 μm sections, which were further picked up onto Superfrost plus microscope slides (Fisherbrand). Slides were positioned at a vertical rack for air dry overnight, placed in the warmer at 55°C for 25 min, and dewaxed for 15 min. Next, the slides were steamed in the citrate buffer for 30 min for antigen retrieval, cooled down on the bench for 20min and cleaned in the water holder for 5min. The tissues were further edged by the ImmEdge pen (Vector Laboratories, Burlingame, CA), blocked with the protein block (Dako, Carpinteria, CA) for 10min at 37°C, incubated with 1:200 diluted primary antibodies at 37°C for 1h, and HRP-conjugated goat-anti-mouse pAb (Jackson ImmunoResearch) as the secondary antibody at 37°C for 1h. The tissue samples were further H&E stained and subjected to BX63 automated fluorescence microscope (Olympus LS) for imaging. We randomly took 7 pictures for each mouse under the 10× and 20× objective lens. The 10× pictures were subjected to the EVOS Analysis software for the automated BMRF1 positive cell counting. In the EVOS Analysis program, five BMRF1 positive cells were selected with the cycle selection tool in the system as the examples of positive signal; five BMRF1 negative cells were selected as the examples of the negative signal. The area threshold was then set between 35-135 to get rid of the cell debris. All the pictures were analyzed with the same setting and exported as a TIF pictures with cell counts on it (Fig. S6C).

## QUANTIFICATION AND STATISTICAL ANALYSIS

### Statistical analysis

Unless otherwise indicated, all bar graphs and line graphs represent the arithmetic mean of three independent experiments (n = 3), with error bars denoting standard deviations. Data were analyzed using two-tailed paired Student t test or analysis of variance (ANOVA) with the appropriate post-test using GraphPad Prism7 software. Gene ontology analysis was done with the Enrichr module using the KEGG pathway databases. Default parameters of Enrichr module was used, with the exception that the Enrichment statistic was set as classic. Metabolic pathway analysis were performed using MetaboAnalyst 3.0.

## Supplemental Figure Legends

**Figure S1. Methionine restriction effects on EBV oncoprotein expression and Burkitt cell growth and survival, related to** Figure 1.

A) Confocal immunofluorescence analysis of LMP1 expression in Mutu I or P3HR-1 cells cultured in media with the indicated methionine concentrations for 3 days. Nuclei were stained by Hoechst. White scale bar indicates 100 µm.

B) Immunoblot analysis of WCL from Mutu I grown for 3 days in RPMI with 10% dialyzed FBS with the indicated methionine concentration. WCL from Mutu III are show at left as a positive control. Representative of n=3 replicates.

C) FACS analysis of PM ICAM1 abundances in the indicated cells cultured for 3 days in RPMI with control 100 μM (blue) vs the indicated methionine restricted concentration (pink).

D) FACS analysis of 7-Aminoactinomycin D (7-AAD) vital dye uptake in P3HR-1 or Mutu I cells cultured in for 3 days. Data are representative of n = 3 biologically independent replicates.

E) Carboxyfluorescein succinimidyl ester (CFSE) cell proliferation analysis of P3HR-1 or Mutu I cells cultured for 3 days in media with 100 μM (blue) vs the indicated methionine restricted concentration (pink). Data are representative of n = 3 replicates.

F) Propidium iodide (PI) cell cycle analysis of P3HR-1 or Mutu I cells cultured for 3 days in media with 100 μM (C) vs methionine restriction (MR, 10 μM for P3HR-1, 1 μM for Mutu I). Data are representative of n = 3 biologically independent replicates. Average percentages from n=3 replicates, as in E.

**Figure S2. Methionine restriction effects on Burkitt EBV gene expression, related to** Figure 1.

A) RNAseq analysis of EBV gene abundances in P3HR-1 (left) or Rael (right) cells cultured in media with 100 μM or 10 μM methionine for 3 days. Heatmaps depict Z-score standard deviation variation from the mean value for each EBV gene across replicates.

B) Scatter plot of log2 fold change of RNAseq values for host and viral mRNAs in Rael (Y-axis) vs P3HR-1 (X-axis) grown in 100 vs 10 μM Met, as in A.

C) Enrichr KEGG pathway –Log 10 (adjusted p-values) of Rael gene sets significantly changed in 100 versus 10 μM methionine, as in (A).

D) Mean ± SEM RNAseq reads of immediate early BZLF1, early BMRF1 or late BLLF1 mRNAs from n = 3 datasets of Rael cells cultured in 100μM versus 10 μM methionine for 3 days. p-values were calculated by unpaired two-sided student’s t-test with equal variance assumption.

E. Immunoblot of WCL from late gene BLLF1 or immediate early gene BZLF1 KO Mutu I cells triggered for lytic replication by anti-IgM crosslinking (1µg/ml).

F. Immunoblot of WCL from BZLF1 KO Mutu I cells cultured for 3 days in media with 100μM versus 10 μM methionine. Blots in E-F are representative of n=3 replicates.

**Figure S3. MR effects on Burkitt methionine cycle and the EBV epigenome. Related to** Figures 1 **and** 2.

A) Mean ± SEM intracellular methionine, SAM, and SAH levels in P3HR-1 cultured for 3 days in media with 100μM versus 10 μM methionine from the experiment presented in Figure 1G.

B) ChIP-qPCR analysis of control IgG, H3K9me3 or H3K27me3 marks at the indicated EBV genomic promoters of Mutu I cells grown in 100μM versus 10 μM methionine for 3 days. Mean ± SEM from n=3 replicates are shown.

C) MeDIP-qPCR analysis of 5mC levels at the indicated EBV genomic promoters of Mutu I cells grown in 100μM versus 10 μM methionine for 3 days. Mean ± SEM from n=3 replicates are shown.

**Figure S4. Burkitt B-cell folate cycle flux is critical for repression of LMP1 target ICAM1 expression. Related to** Figure 5.

A) Mean ± SEM PM ICAM1 values from n=3 replicates of Mutu I cells grown in RPMI with 10% dialyzed or undialyzed FBS and treated with DMSO, SHIN1 (10μM) and/or sodium formate (1mM) as indicated, as in Fig. 5A.

B) FACS analysis of PM ICAM1 abundances in Rael cells grown in RPMI with 10% undialyzed versus dialyzed serum and treated with SHIN1 (10μM) and/or sodium formate (1mM) as indicated.

C) Mean ± SEM plasma membrane ICAM1 values from n=3 replicates of Rael cells as in Fig. (B).

**Figure S5. Dietary MR mouse xenograft pilot studies. Related to** Figure 6.

A) Schematic of dietary MR pilot study. Mutu I tumors were implanted in mouse flanks at two weeks after initiation of control versus MR diet. Samples were collected at week 5.

B) Mass spectrometry analysis of serum methionine values in n=3 mice at day 0 and 7 of the MR diet prior to tumor implantation. Lines connect paired samples from a given mouse.

C) Mean ± SEM body weight measurements for mice on control (black) vs MR (red) diets.

D) Mean ± SEM Mutu I tumor volumes in mice fed control vs MR diets.

**Figure S6. Dietary MR effects on Mutu I xenograft methionine cycle and BMRF1 expression. Related to** Figure 6.

A) Mean ± SEM levels of the indicated tumor and plasma methionine cycle component from n=3 samples drawn at day 14 of the control or MR diet.

B) Mean ± SEM body weight measurements for mice on control (gray) vs MR (blue) diets.

C) Immunohistochemical analysis of BMRF1 expression in two representative tumor fields from mice fed control or MR diets for 14 days.

**Figure S7. SHIN2 effects on Mutu I xenograft BMRF1 expression. Related to** Figure 6.

A) Immunohistochemical analysis of BMRF1 expression in two representative tumor fields from mice treated with vehicle control or SHIN2 3 days prior, as in Figure 6I.

B) Mean ± SEM numbers of BMRF1+ cells per 10X field as in (A).

