## Supplemental Figures for "Methionine Metabolism Controls the B-cell EBV Epigenome and Viral Latency"

**Fig. S1**

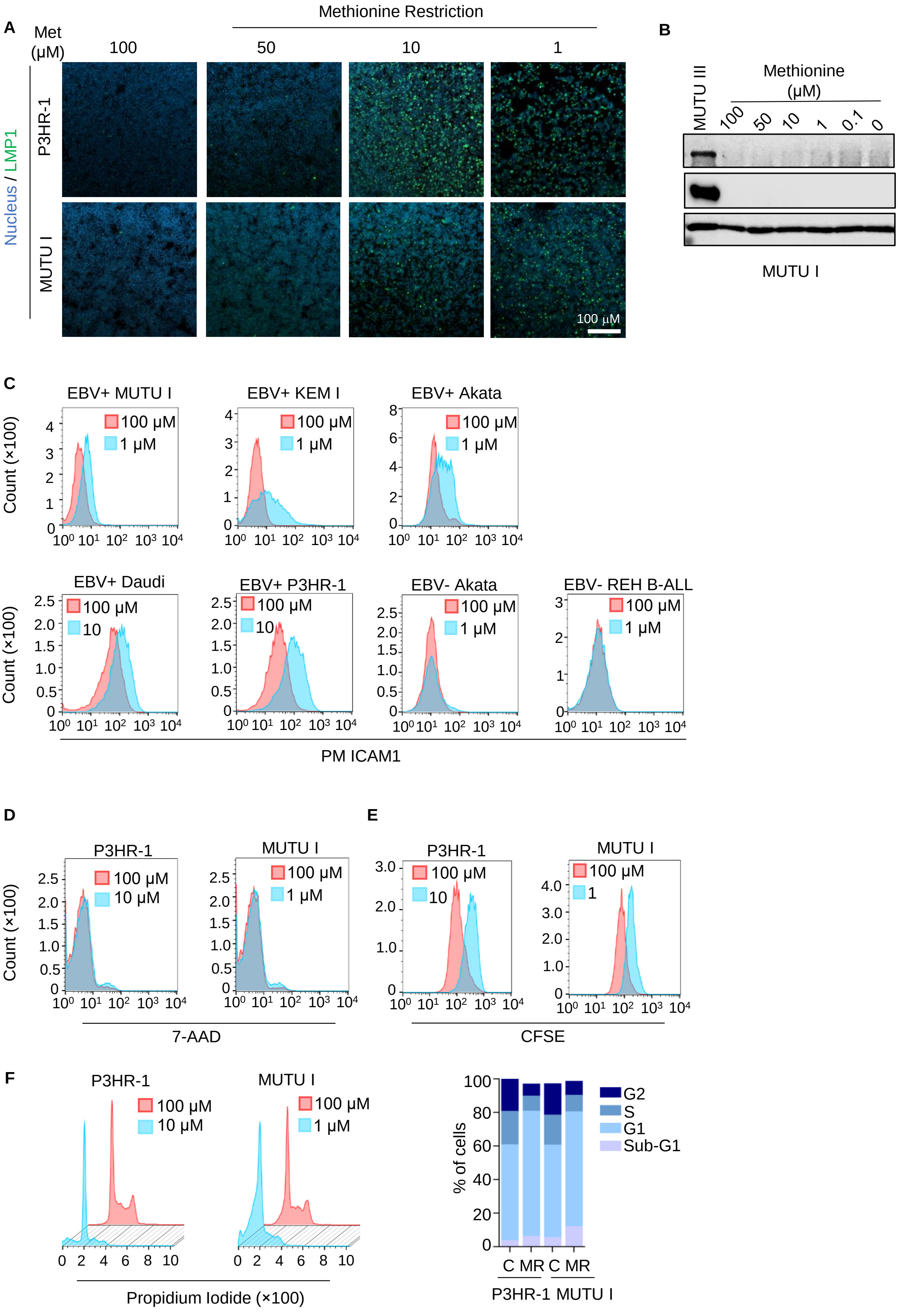
A
Methionine Restriction
Met (μM)
100
50
10
1
Nucleus / LMP1
P3HR-1
MUTU I
100 μM
B
MUTU III
Methionine (μM)
100
50
10
1
0.1
0
EBNA3C
EBNA2
GAPDH
MUTU I
C
EBV+ MUTU I
EBV+ KEM I
EBV+ Akata
EBV+ Daudi
EBV+ P3HR-1
EBV- Akata
EBV- REH B-ALL
100 μM
1 μM
10
Count (×100)
PM ICAM1
D
P3HR-1
MUTU I
100 μM
10 μM
1 μM
Count (×100)
7-AAD
E
P3HR-1
MUTU I
100 μM
10
1
CFSE
F
P3HR-1
MUTU I
100 μM
10 μM
1 μM
Propidium Iodide (×100)
% of cells
G2
S
G1
Sub-G1
C
MR
C
MR
P3HR-1
MUTU I

**Figure S1. Methionine restriction effects on EBV oncoprotein expression and Burkitt cell growth and survival, related to Figure 1.**

A) Confocal immunofluorescence analysis of LMP1 expression in Mutu I or P3HR-1 cells cultured in media with the indicated methionine concentrations for 3 days. Nuclei were stained by Hoechst. White scale bar indicates 100 µm.

B) Immunoblot analysis of WCL from Mutu I grown for 3 days in RPMI with 10% dialyzed FBS with the indicated methionine concentration. WCL from Mutu III are show at left as a positive control. Representative of n=3 replicates.

C) FACS analysis of PM ICAM1 abundances in the indicated cells cultured for 3 days in RPMI with control 100 µM (blue) vs the indicated methionine restricted concentration (pink).

D) FACS analysis of 7-Aminoactinomycin D (7-AAD) vital dye uptake in P3HR-1 or MUTU I cells cultured in for 3 days. Data are representative of n = 3 biologically independent replicates.

E) Carboxyfluorescein succinimidyl ester (CFSE) cell proliferation analysis of P3HR-1 or Mutu I cells cultured for 3 days in media with 100 µM (blue) vs the indicated methionine restricted concentration (pink). Data are representative of n = 3 replicates.

F) Propidium iodide (PI) cell cycle analysis of P3HR-1 or Mutu I cells cultured for 3 days in media with 100 µM (C) vs methionine restriction (MR, 10 µM for P3HR-1, 1 µM for MUTU I). Data are representative of n = 3 biologically independent replicates. Average percentages from n=3 replicates, as in E.

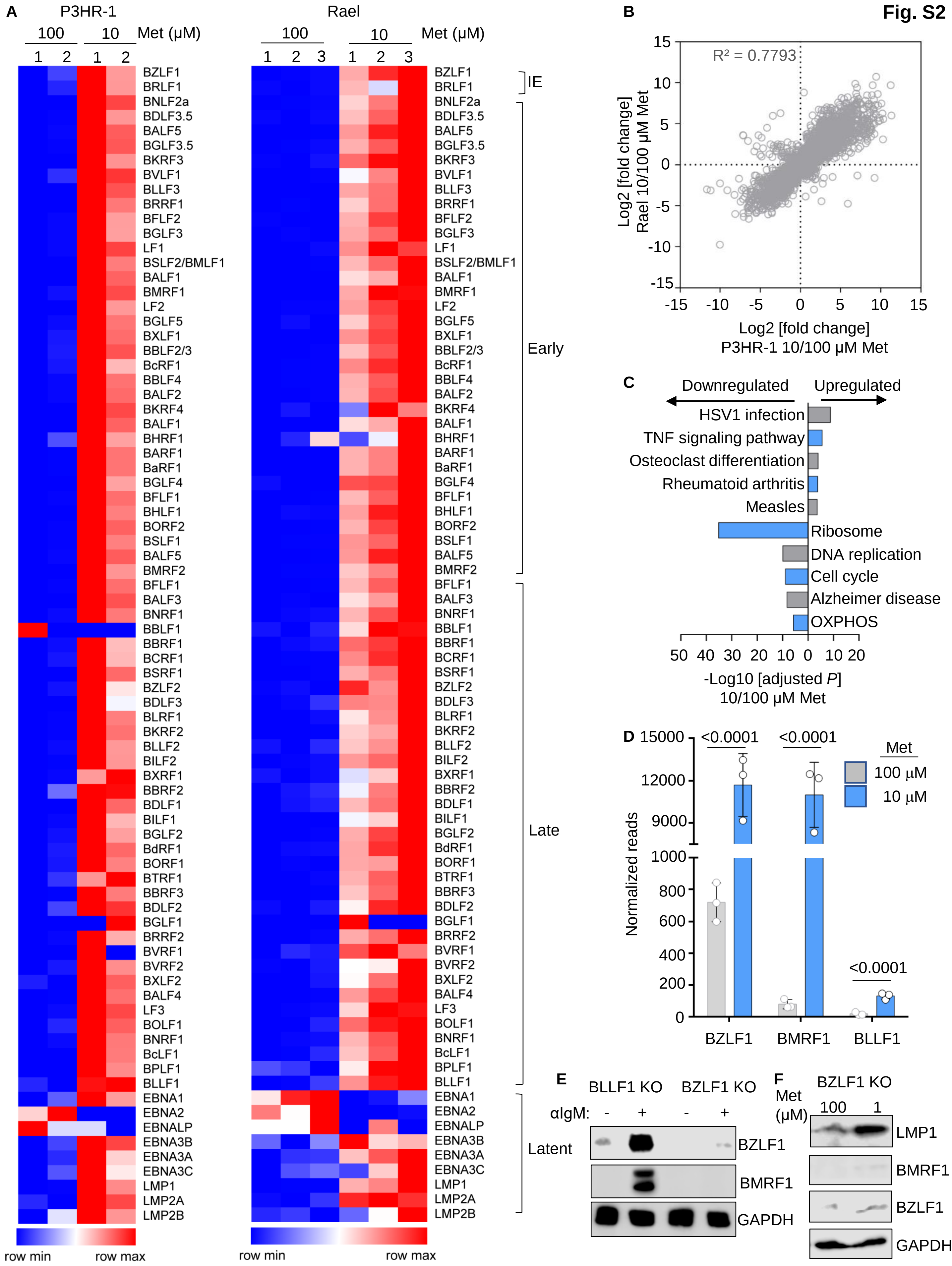

Fig. S2
A
P3HR-1
Rael
Met (μM)
100
10
IE
Early
Late
Latent
BZLF1
BRLF1
BNLF2a
BDLF3.5
BALF5
BGLF3.5
BKRF3
BVLF1
BLLF3
BRRF1
BFLF2
BGLF3
LF1
BSLF2/BMLF1
BALF1
BMRF1
LF2
BGLF5
BXLF1
BBLF2/3
BcRF1
BBLF4
BALF2
BKRF4
BHRF1
BARF1
BaRF1
BGLF4
BFLF1
BHLF1
BORF2
BSLF1
BMRF2
BALF3
BNRF1
BBLF1
BBRF1
BCRF1
BSRF1
BZLF2
BDLF3
BLRF1
BKRF2
BLLF2
BILF2
BXRF1
BBRF2
BDLF1
BILF1
BGLF2
BdRF1
BORF1
BTRF1
BBRF3
BDLF2
BGLF1
BRRF2
BVRF1
BVRF2
BXLF2
BALF4
LF3
BOLF1
BcLF1
BPLF1
BLLF1
EBNA1
EBNA2
EBNALP
EBNA3B
EBNA3A
EBNA3C
LMP1
LMP2A
LMP2B
row min
row max
B
R² = 0.7793
Log2 [fold change]
Rael 10/100 μM Met
Log2 [fold change]
P3HR-1 10/100 μM Met
C
Downregulated
Upregulated
HSV1 infection
TNF signaling pathway
Osteoclast differentiation
Rheumatoid arthritis
Measles
Ribosome
DNA replication
Cell cycle
Alzheimer disease
OXPHOS
-Log10 [adjusted P]
10/100 μM Met
D
Normalized reads
<0.0001
Met
100 μM
10 μM
BZLF1
BMRF1
BLLF1
E
BLLF1 KO
BZLF1 KO
αIgM:
BZLF1
BMRF1
GAPDH
F
BZLF1 KO
Met (μM)
LMP1
BMRF1
BZLF1
GAPDH

**Figure S2. Methionine restriction effects on Burkitt EBV gene expression, related to Figure 1.**

A) RNAseq analysis of EBV gene abundances in P3HR-1 (left) or Rael (right) cells cultured in media with 100 μM or 10 μM methionine for 3 days. Heatmaps depict Z-score standard deviation variation from the mean value for each EBV gene across replicates.

B) Scatter plot of log2 fold change of RNAseq values for host and viral mRNAs in Rael (Y-axis) vs P3HR-1 (X-axis) grown in 100 vs 10 μM Met, as in A.

C) Enrichr KEGG pathway –Log 10 (adjusted p-values) of Rael gene sets significantly changed in 100 versus 10 μM methionine, as in (A).

D) Mean ± SEM RNAseq reads of immediate early BZLF1, early BMRF1 or late BLLF1 mRNAs from n = 3 datasets of Rael cells cultured in 100μM versus 10 μM methionine for 3 days. p-values were calculated by unpaired two-sided student's t-test with equal variance assumption.

E. Immunoblot of WCL from late gene BLLF1 or immediate early gene BZLF1 KO Mutu I cells triggered for lytic replication by anti-IgM crosslinking (1μg/ml).

F. Immunoblot of WCL from BZLF1 KO Mutu I cells cultured for 3 days in media with 100μM versus 10 μM methionine. Blots in E-F are representative of n=3 replicates.

Fig. S3

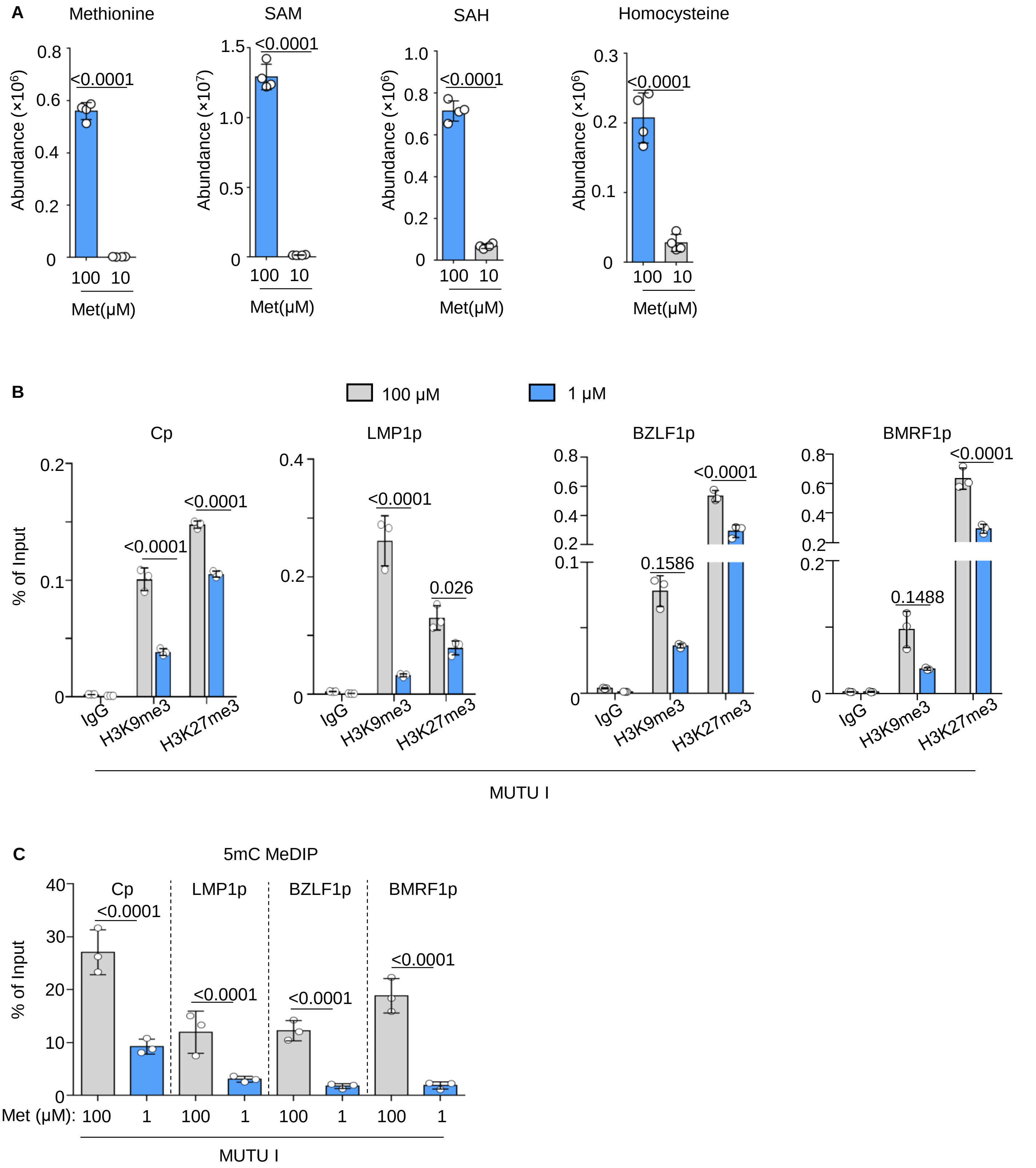
A
Methionine
SAM
SAH
Homocysteine
Abundance (×10^6)
Abundance (×10^7)
<0.0001
Met(μM)
100
10
B
100 μM
1 μM
Cp
LMP1p
BZLF1p
BMRF1p
% of Input
<0.0001
0.026
0.1586
0.1488
IgG
H3K9me3
H3K27me3
MUTU I
C
5mC MeDIP
Cp
LMP1p
BZLF1p
BMRF1p
% of Input
<0.0001
Met (μM):
100
1
MUTU I

**Figure S3. MR effects on Burkitt methionine cycle and the EBV epigenome. Related to Figures 1 and 2.**

A) Mean ± SEM intracellular methionine, SAM, and SAH levels in P3HR-1 cultured for 3 days in media with 100µM versus 10 µM methionine from the experiment presented in Figure 1G.

B) ChIP-qPCR analysis of control IgG, H3K9me3 or H3K27me3 marks at the indicated EBV genomic promoters of Mutu I cells grown in 100µM versus 10 µM methionine for 3 days. Mean ± SEM from n=3 replicates are shown.

C) MeDIP-qPCR analysis of 5mC levels at the indicated EBV genomic promoters of MUTU I cells grown in 100µM versus 10 µM methionine for 3 days. Mean ± SEM from n=3 replicates are shown.

Fig.S4

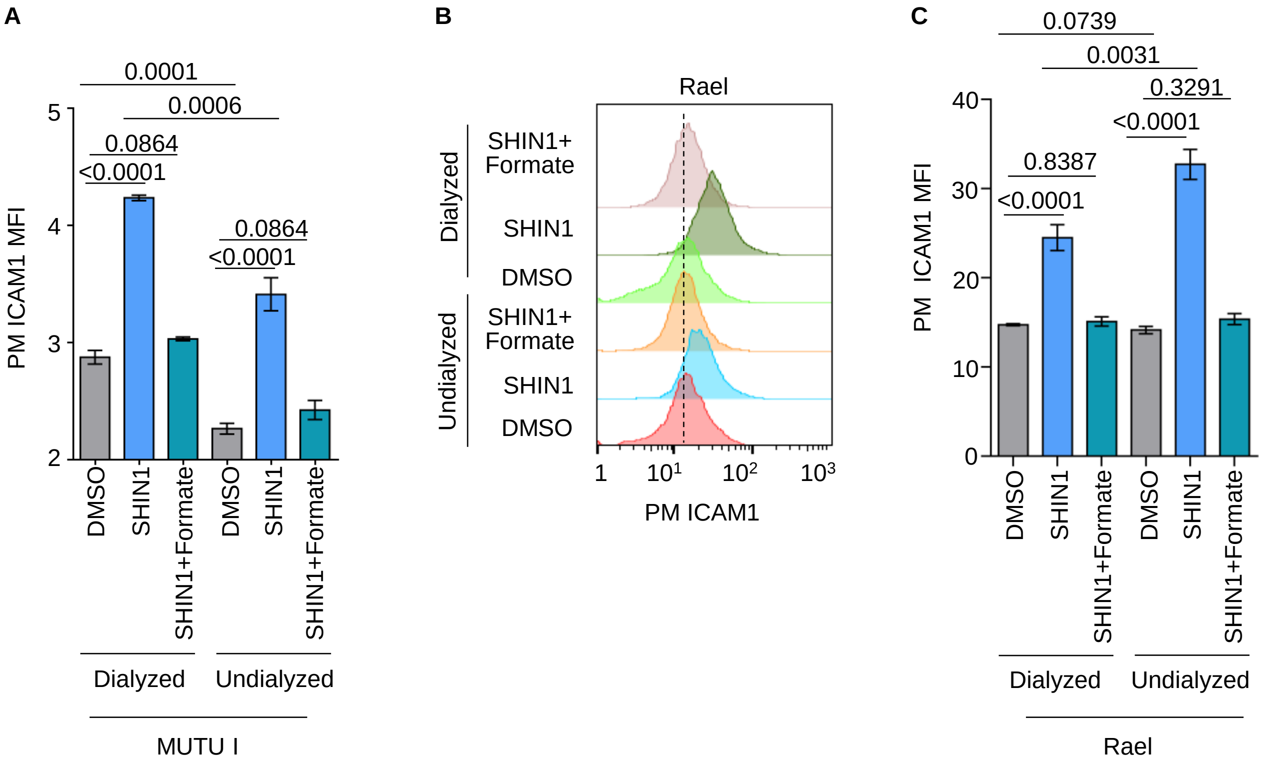
A
0.0001
0.0006
0.0864
<0.0001
0.0864
<0.0001
PM ICAM1 MFI
5
4
3
2
DMSO
SHIN1
SHIN1+Formate
DMSO
SHIN1
SHIN1+Formate
Dialyzed
Undialyzed
MUTU I
B
Rael
Dialyzed
Undialyzed
SHIN1+ Formate
SHIN1
DMSO
SHIN1+ Formate
SHIN1
DMSO
1
10^1
10^2
10^3
PM ICAM1
C
0.0739
0.0031
0.3291
<0.0001
0.8387
<0.0001
PM ICAM1 MFI
40
30
20
10
0
DMSO
SHIN1
SHIN1+Formate
DMSO
SHIN1
SHIN1+Formate
Dialyzed
Undialyzed
Rael

**Figure S4. Burkitt B-cell folate cycle flux is critical for repression of LMP1 target ICAM1 expression. Related to Figure 5.**

A) Mean ± SEM PM ICAM1 values from n=3 replicates of Mutu I cells grown in RPMI with 10% dialyzed or undialyzed FBS and treated with DMSO, SHIN1 (10μM) and/or sodium formate (1mM) as indicated, as in Fig. 5A.

B) FACS analysis of PM ICAM1 abundances in Rael cells grown in RPMI with 10% undialyzed versus dialyzed serum and treated with SHIN1 (10μM) and/or sodium formate (1mM) as indicated.

C) Mean ± SEM plasma membrane ICAM1 values from n=3 replicates of Rael cells as in Fig. (B).

Fig.S5

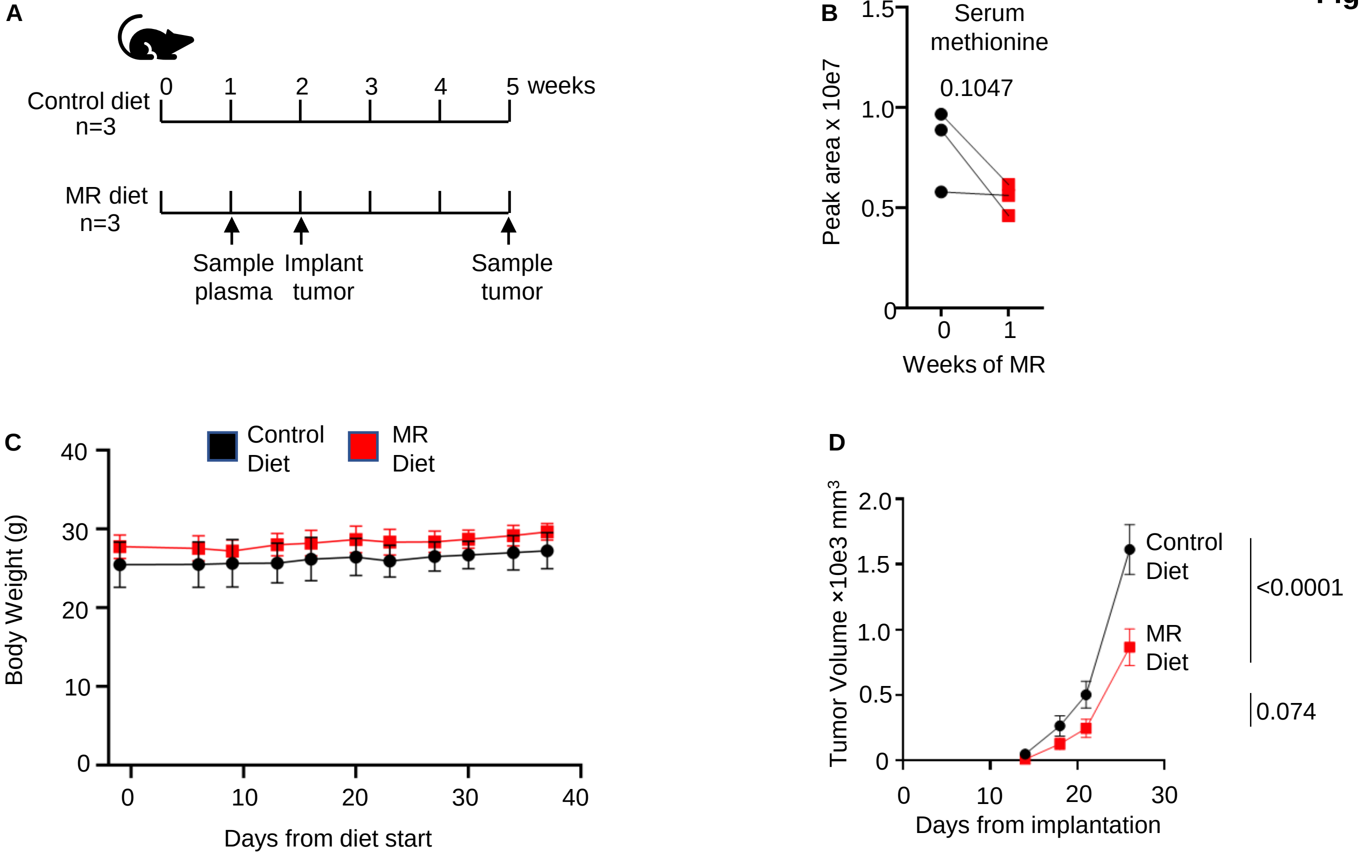

A
Control diet
n=3
0
1
2
3
4
5 weeks
MR diet
n=3
Sample plasma
Implant tumor
Sample tumor
B
Serum methionine
0.1047
Peak area x 10e7
1.5
1.0
0.5
0
0
1
Weeks of MR
C
Control Diet
MR Diet
Body Weight (g)
40
30
20
10
0
0
10
20
30
40
Days from diet start
D
Tumor Volume ×10e3 mm³
2.0
1.5
1.0
0.5
0
Control Diet
MR Diet
<0.0001
0.074
0
10
20
30
Days from implantation

**Figure S5. Dietary MR mouse xenograft pilot studies. Related to Figure 6.**

A) Schematic of dietary MR pilot study. Mutu I tumors were implanted in mouse flanks at two weeks after initiation of control versus MR diet. Samples were collected at week 5.

B) Mass spectrometry analysis of serum methionine values in n=3 mice at day 0 and 7 of the MR diet prior to tumor implantation. Lines connect paired samples from a given mouse.

C) Mean ± SEM body weight measurements for mice on control (black) vs MR (red) diets.

D) Mean ± SEM Mutu I tumor volumes in mice fed control vs MR diets.

**Fig.S6**

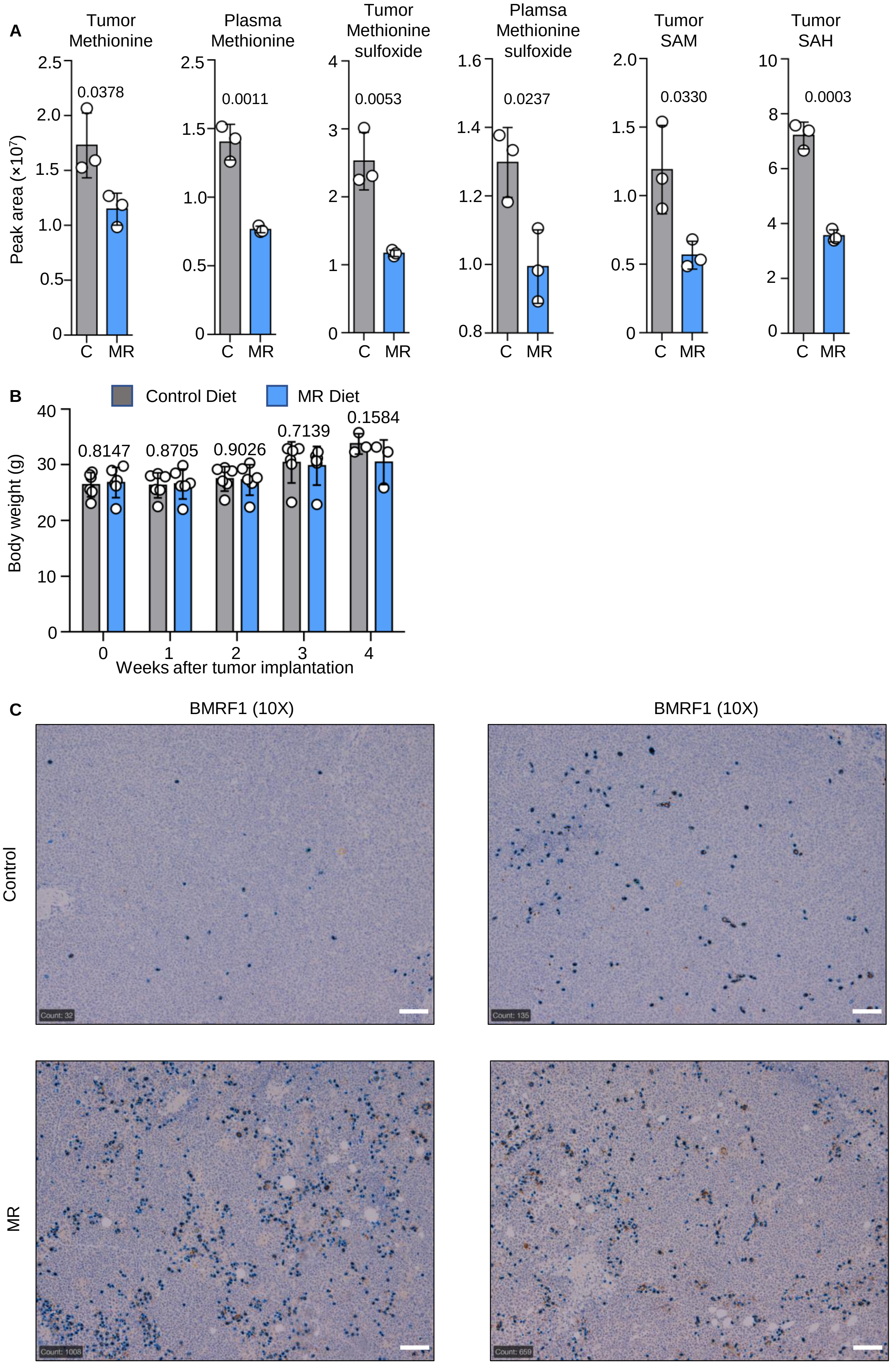
A
Tumor Methionine
Plasma Methionine
Tumor Methionine sulfoxide
Plamsa Methionine sulfoxide
Tumor SAM
Tumor SAH
Peak area (×10^7)
0.0378
0.0011
0.0053
0.0237
0.0330
0.0003
C
MR
B
Control Diet
MR Diet
Body weight (g)
0.8147
0.8705
0.9026
0.7139
0.1584
Weeks after tumor implantation
C
BMRF1 (10X)
BMRF1 (10X)
Control
MR
Count: 32
Count: 135
Count: 1008
Count: 659

**Figure S6. Dietary MR effects on Mutu I xenograft methionine cycle and BMRF1 expression. Related to Figure 6.**

A) Mean ± SEM levels of the indicated tumor and plasma methionine cycle component from n=3 samples drawn at day 14 of the control or MR diet.

B) Mean ± SEM body weight measurements for mice on control (gray) vs MR (blue) diets.

C) Immunohistochemical analysis of BMRF1 expression in two representative tumor fields from mice fed control or MR diets for 14 days. Scale bar, 100nm.

**Fig. S7**

**A**

BMRF1 10× | BMRF1 20×

Control

SHIN2

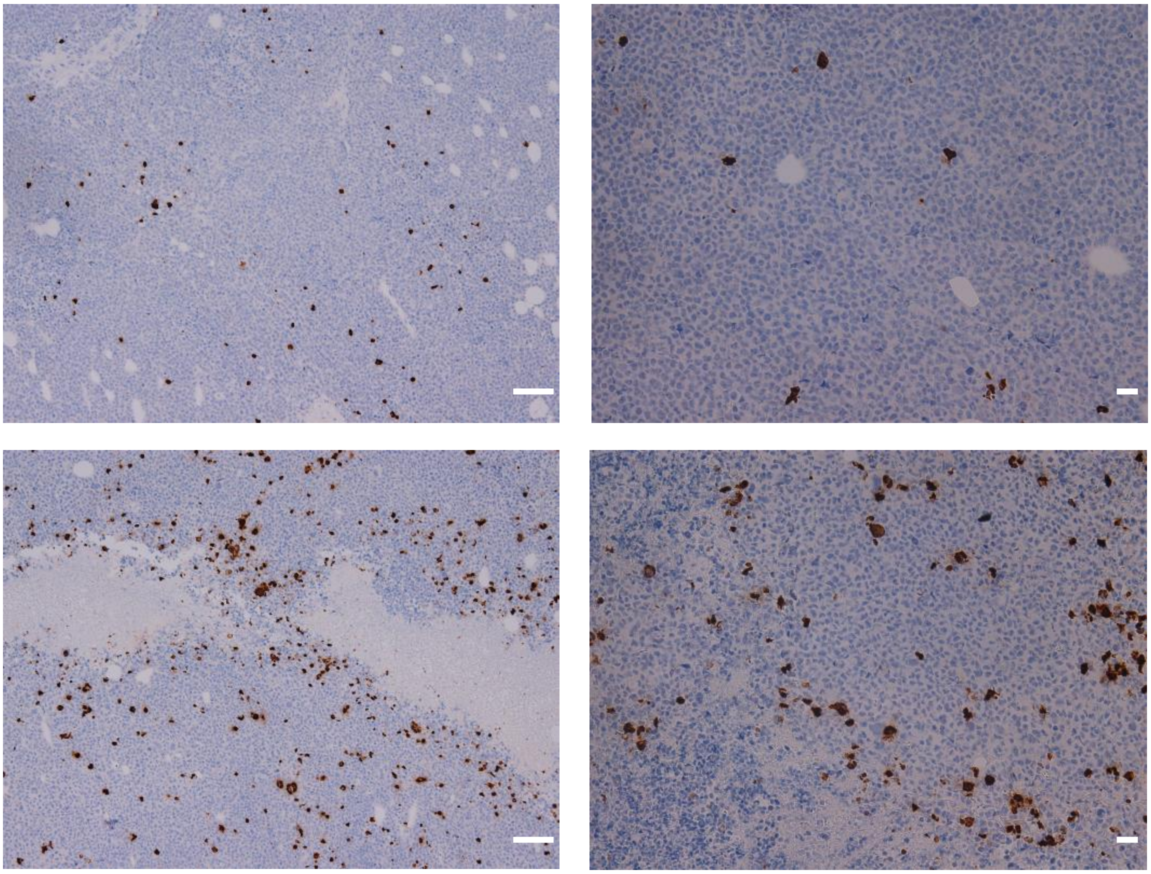

Day 3

**B**

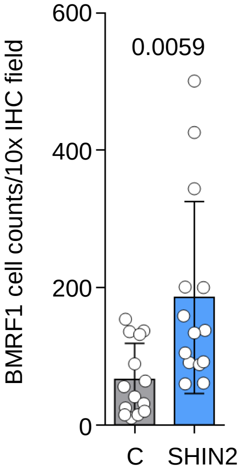

**Figure S7. SHIN2 effects on Mutu I xenograft BMRF1 expression. Related to Figure 6.**

A) Immunohistochemical analysis of BMRF1 expression in two representative tumor fields from mice treated with vehicle control or SHIN2 3 days prior, as in **Figure 6I**. Scale bar, 100nm.

B) Mean ± SEM numbers of BMRF1+ cells per 10X field as in (A).
